# Domestication and lowland adaptation of coastal preceramic maize from Paredones, Peru

**DOI:** 10.1101/2022.02.23.481166

**Authors:** Miguel Vallebueno-Estrada, Guillermo G. Hernández-Robles, Eduardo González-Orozco, Iván López-Valdivia, Teresa Rosales Tham, Víctor Vásquez Sánchez, Kelly Swarts, Tom D. Dillehay, Jean-Philippe Vielle-Calzada, Rafael Montiel

**Affiliations:** Grupo de Desarrollo Reproductivo y Apomixis, Unidad de Genómica Avanzada, Laboratorio Nacional de Genómica para la Biodiversidad, CINVESTAV Irapuato, México; Grupo de Interacción Núcleo-Mitocondrial y Paleogenómica, Unidad de Genómica Avanzada, Laboratorio Nacional de Genómica para la Biodiversidad, CINVESTAV Irapuato, México; Departamento de Antropología, Universidad Nacional de Trujillo, Perú; Centro de Investigaciones Arquebiológicas y Paleoecológicas Andinas ARQUEBIOS, Apartado Postal 595, Trujillo, Perú; Max Planck Institute for Developmental Biology. 72076 Tübingen, Germany; Department of Anthropology, Vanderbilt University, Nashville TN 37235; Escuela de Arqueología, Universidad Austral de Chile, Puerto Montt, Chile

**Keywords:** maize, paleogenomics, domestication, Paredones, highlands

## Abstract

Archaeological cobs from Paredones and Huaca Prieta (Peru) are phenotypically indistinguishable from modern maize. This contrasts with the earliest Mexican macro-specimens from Guila Naquitz and San Marcos, which are phenotypically intermediate even though they date more recently in time. These observations suggest at least two alternative scenarios, one in which maize was domesticated earlier than previously thought in the lowland Mesoamerica, followed by rapid lowland dispersal to Peru, and another in which maize was independently domesticated in South America and subsequently lost, as current evidence supports a single origin for all modern maize. To gain insights into the origins of ancient Peruvian maize, we sequenced DNA from three Paredones specimens dating 6775 to 5000 calibrated years before present (BP) and conducted comparative analyses with two teosinte subspecies (*Zea mays* ssp. *mexicana* and *parviglumis*) and extant maize, including highland and lowland landraces from Mesoamerica and South America. We show that Paredones maize originated from the same domestication event as Mexican maize and was domesticated by 6775 BP, implying rapid dispersal followed by improvement. Paredones maize show minimal levels of gene flow from *mexicana*, smaller than those observed in teosinte *parviglumis*. It also harbors significantly fewer alleles previously found to be adaptive to highlands, but not of alleles adaptive to lowlands, supporting a lowland migration route. Our overall results imply that Paredones maize originated in Mesoamerica, arrived in Peru without *mexicana* introgression through a rapid lowland migration route, and underwent improvements in both Mesoamerica and South America.

**Significance Statement:** The coastal Peruvian preceramic sites of Paredones and Huaca Prieta provide the earliest known maize macro-remains. Found more than 3,800 km away from the maize center of origin and presenting a phenotypically modern cob constitution relative to their antiquity, these specimens represent a paradox for understanding maize evolution and dispersal. We show that Paredones maize originated in Mesoamerica, like all known maize, and arrived in South America without introgression from the teosinte *mexicana*. Since modern maize has substantial contributions from *mexicana*, it raises the question of when *mexicana* introgression spread to South America. Paredones maize preferentially shares adaptive allelic diversity with lowland Mesoamerican samples, suggesting a migration route probably associated with a coastal corridor previously identified with archeological findings.

## Introduction

Although the elucidation of the origins of maize (*Zea mays* ssp. *mays L*.) based on archeological data puzzled the scientific community for several decades (1–3), the integration of genomic, archeological, and botanical evidence has identified the Balsas basin in central Mexico as the only center of origin of maize (4–6), and that the divergence from its wild ancestor, *Zea mays* ssp. *parviglumis* (hereafter *parviglumis*), occurred about 9,000 years ago (4, 7). Domestication occurred in a single event creating a monophyletic clade that includes all domesticated maize landraces and diverges from both *parviglumis* and *Zea mays* ssp. *mexicana* (hereafter *mexicana*) populations (4). Genomic investigations of archeological samples from the Tehuacan highland site suggested that the dispersal of maize to the highlands of México was complex, as early-arriving maize populations retained higher levels of genomic diversity than expected (8, 9). The constant gene flow between domesticated maize with already divergent populations of *parviglumis* and *mexicana* has contributed to the adaptation of maize to new environments and remains embedded in the genetic structure of its populations (10, 11). Geographic areas of contact have been stable over time, as these teosinte populations have maintained a discrete distribution in central Mexico since the last glacial maximum (12). Gene flow from a sympatric *mexicana* population to domesticated maize populations has been associated with an altitudinal cline in the highlands of Mexico and Guatemala (11, 13, 14), and the genetic introgression from *mexicana* in the form of distinct chromosomal inversions has been associated with adaptation of maize to central Mexican highlands (13, 14). The genetic contribution of teosinte *mexicana* to Mexican highland landraces is about 20% (11, 13), and the time of the introgression from *mexicana* seems to be around 1000 generations ago (15).

Archaeological evidence supports the dispersal of maize populations associated with a Pacific lowland coastal corridor (16–18). Population substructure and differentiation patterns suggested independent adaptations to highland environments in Mesoamerica and South America; meanwhile, minimal population sub-structuring was detected between the lowlands of Mesoamerica and South America (10, 19). While the introgression from *mexicana* of large chromosomal inversions located on chromosomes 3, 4 and 6 has been shown to contribute to adaptation to Mesoamerican highlands (20), those regions were not detected in highland maize populations of South America (14) or North America (10) which were also isolated from direct gene flow with *parviglumis* (21). Although these inversions are specific to Mexican landraces, many maize populations across the Americas including South America show genome wide admixture with *mexicana* (10). Highland South American landraces also show phenotypic diversity relative to lowlands (22), as well as specific cytogenetic characteristics such as the absence of supernumerary highly heterochromatic B chromosomes in Peruvian landraces, resulting in the so-called Andean Complex (23). These unique characteristics have puzzled the scientific community regarding the origin and adaptation of the Andean Complexes (16, 22–26). The archeological expeditions in the coastal Peruvian sites of Paredones and Huaca Prieta yielded a robust collection of ancient maize remains that provide a unique opportunity to investigate the chronology, landrace evolution, and cultural context associated with early maize dispersal in South America (17, 27, 28). These findings include one charred cob fragment dated 6775 to 6504 calibratedBP, and other burned and unburned cobs stratigraphically dated to similar age, representing the most ancient maize macro-specimens found to date. Strikingly, and in contrast to Mexican cob fragments from Guilá Naquitz dating approximately ∼300 years younger (6235 calibrated BP) (29), the Paredones and Huaca Prieta specimens are robust, slender and cylindrical 2.4-3.1 cm long cobs, with eight rows of kernels consistent with the hypothetical Proto-Confite Morocho landrace (28, 29). Given its early presence in the region and its advanced phenotype, some authors considered the possibility of independent domestication for maize in South America (22, 27). Recently, it has been proposed that the first maize lineages arriving in South America were partial domesticates, locally evolving the full set of domestication traits due to reduced gene flow from wild relatives that enhanced anthropogenic pressures (21, 30). However, it is not clear how fast this process could have been and if the earliest archeological samples found in South America were partially or fully domesticated. In addition, the expectation on the phenotype of those hypothetical samples is not clear.

Here we present the genomic analysis of three ancient specimens belonging to the earliest cultural phase of Paredones and dating 6775 to 6504, 5800 to 5400, and 5583 to 5324 calibrated years BP. To reveal the population context of their origin and domestication, we conducted comparisons with *parviglumis, mexicana* and extant maize landraces. Also, to explore if these ancient maize samples exhibit some evidence of *mexicana* gene flow, we performed D-statistics under several experimental designs, comparing them to extant maize and *parviglumis* populations. Finally, we did a comparison with previously published data from extant highland and lowland Mesoamerican and South American landraces, to identify signatures of specific adaptation that could bring insights into the specific improvements that this maize went through in both Mesoamerica and South America. Our results provide evidence that ancient Peruvian maize originated in Mesoamerica as all landraces found to date, followed by a rapid dispersal into the lowlands of South America, and subsequently subjected to local adaptation processes.

## Results

### Paredones ancient maize sampling

The maize macroremains were collected as part of published excavations at the Paredones and Huaca Prieta sites (27). Macroremains from both sites were excavated in deeply stratified and undisturbed cultural floors. Stratigraphic Unit 22 at Paredones is the archeological component with the largest and most diversified amount of maize remains, with the oldest ^14^C dated cobs. The oldest cobs derive from the base of this unit, in a single, discrete and intact floor of ∼2 cm in thickness and at 5.5 m in depth from the present-day surface (17). The dated remains at both sites are chrono-stratigraphically bracketed by and in agreement with more than 160 dates from mound and off-mound contexts that were obtained by Accelerator Mass Spectrometry (AMS) and Optically Stimulated Luminescence (OSL) (17). No taphonomic or other disturbing cultural or geological features were observed in any excavation units that would have altered the integrity and intactness of strata containing the maize remains (*SI Appendix*, Material and Methods). All radiocarbon-dated remains were assayed by the SHCal04 Southern Hemisphere Calibration 0-11.0 calibrated kyr BP curve (31).

In 2019, Dillehay and geologists Steven Goodbred and Elizabeth Chamberlain carried out excavations in a Preceramic domestic site (S-18) located ∼3.2 km north of Huaca Prieta. Preceramic corn remains were encountered consistently in the upper to lower intact cultural layers of the site. As with the Paredones and Huaca Prieta sites, the lower strata contained both charred and uncharred cobs 2.6-3.1 cm long, slender and cylindrical with eight rows of kernels, of the smaller and earliest type of identified corn species Proto Confite Morocho (27) (Grobman, personal communication, 2019). The middle to upper strata yielded the known later and slightly larger Preceramic varieties of Confite Chavinese and Proto Alazan. An OSL date from a discrete and intact lower layer containing a hearth with two unburned cob fragments of the Proto Confite Morocho variety assayed ∼7000 +/-630 years ago or 5610-4350 BCE (32). Wood charcoal from the hearth was processed at 7162-6914 +/-30calibratedl BP (AA75398), indicating that the associated cob fragments date ∼7000 years ago. In South America, maize micro remains (e.g. starch grains, pollen, phytoliths) have been dated ∼7500–7000 calibrated BP (33–35) at sites in southwest coastal Ecuador, located ∼450 km north of Huaca Prieta and Paredones, and in other localities across the continent at ∼6500 calibrated BP and later (21).

Excavation Units 20 and 22 from Paredones are illustrated in Figs. 1A and 1B. Three of the recovered maize samples (Par_N1, Par_N9, and Par_N16) are well-structured maize cobs deprived of seeds and showing morphological similarities to extant landraces. Par_N1 is the most ancient specimen, dating 5,900±40 ^14^C years BP (6775-6504 2σ calibrated BP at 95% confidence), and obtained from archeological Unit 22 (Fig. 1C). The other two samples found in Unit 20, Par_N9 and Par_N16, were dated 5800–5400 and 5583-5324 calibrated BP, respectively (Fig. 1C and *SI Appendix*, Table S1). Par_N1 is older than any other maize macro-specimen found to date (36).

**Figure 1.**
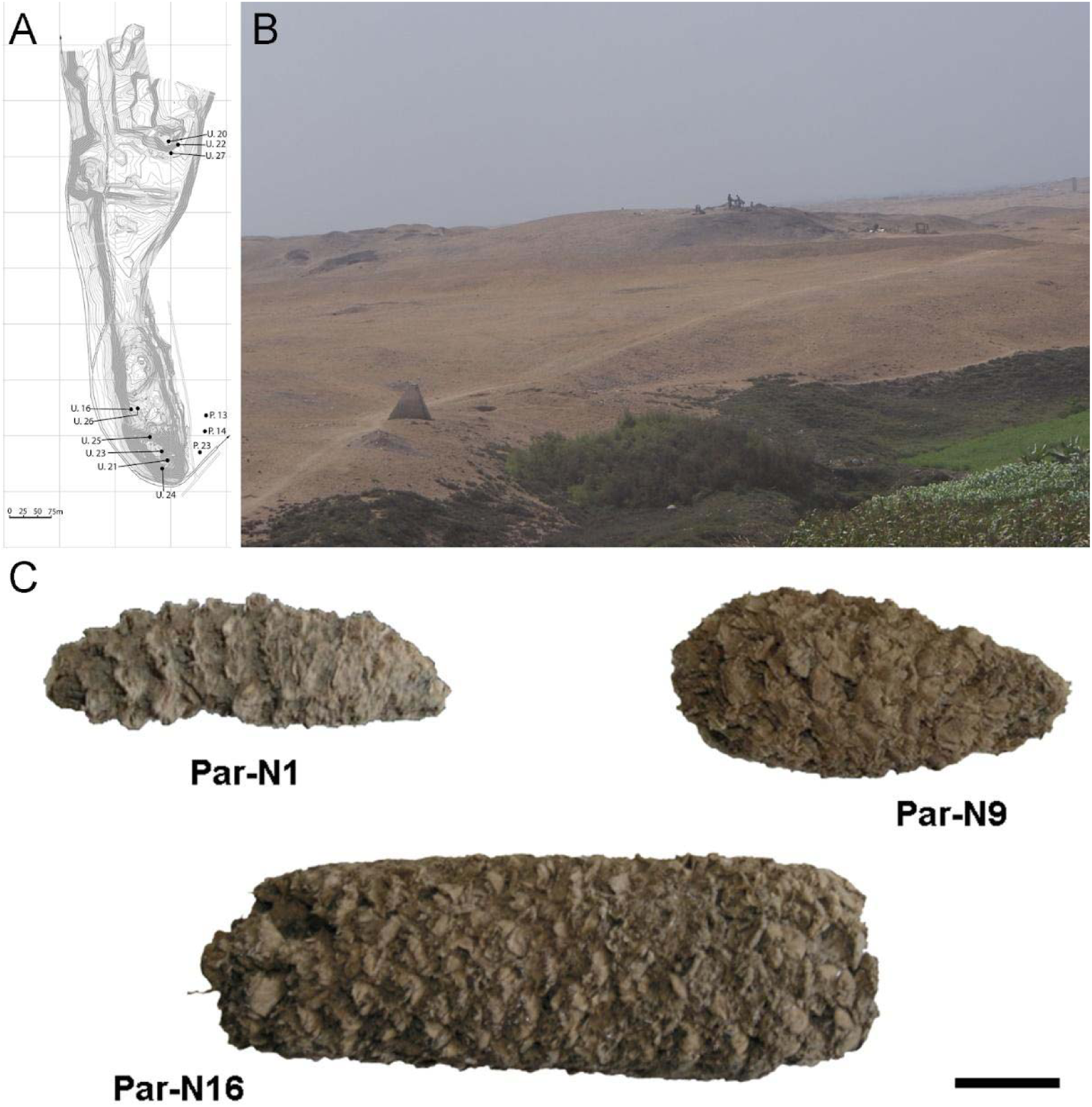
Archeological site and specimens of Paredones. (**A**) Topographic contour map of Huaca Prieta and Paredones (units U20, U22, and U27) coastal sites, showing excavation units. (**B**) The Paredones mound during archeological excavations. (**C**) Maize specimens Par_N1 (dating 6,775-6,504 calibrated years BP), Par_N9 (dating to 5,800–5,400 calibrated years BP), and ParN16 (dating 5,583-5,324 calibrated years BP); Scale bar =1 cm.

### Paleogenomic characterization of ancient maize samples

To determine the genomic constitution and degree of genetic variability present in the 6775-5324 BP maize of Paredones, we extracted DNA and conducted whole-genome shotgun sequencing in specimens Par_N1, Par_N9, and Par_N16. Since the endogenous DNA content of all three specimens was low (0.2% for Par_N1 and Par_N9; 1.1% for Par_N16), we conducted in-depth whole-genome shotgun sequencing of high-quality libraries under Illumina platforms, generating 622 million (M) quality-filtered reads for Par_N1, 423M for Par_N9, and 392M for Par_N16. Further sequencing of the Par_N16 library yielded 459M additional reads, to generate a total of 851M for this sample (*SI Appendix*, Table S2). Comparison with version 3 of the B73 maize reference genome resulted in 1,320,284 (Par_N1), 1,034,544 (Par_N9), and 15,023,803 (Par_N16) reads mapping to either repetitive (33.4% for Par_N1; 34% for Par_N9; 34.8% for Par_N16) or unique (66.5% for Par_N1; 66% for Par_N9; 65.2% for Par_N16) genomic regions, for a total virtual length of 52.2 Mb (Par_N1), 40.8 Mb (Par_N9), and 471.65 Mb (Par_N16) of the unique maize genome (SI Appendix, Table S2). Average mapping quality in Phred score was 31.9 for Par_N1, 32.1 for Par_N9 and 34 for Par_N16; this is reflected in the estimated error rate of 1.19E-02 for Par_N1, 9.59E-03 for Par_N9 and 1.05E-02 for Par_N16 (*SI Appendix*, Table S2). Reads contained signatures of DNA damage typical of postmortem degradation in ancient samples, including overhangs of single-stranded DNA, 13-20% cytosine deamination and fragmentation due to depurination (37) resulting in median fragment lengths of 36 bp for all three samples. A total of 42% to 53% of all covered sites had signatures of molecular damage (*SI Appendix*, Fig. S1). This damage pattern is an indication that this is ancient endogenous DNA and does not represent DNA contamination from extant sources. After mapping reads corresponding to unique genomic regions, Par_N1, Par_N9 and Par_N16 yielded approximately 16.9M (Par_N1), 12.1M (Par_N9), and 334.36M (Par_N16) unique genomic sites spread across all 10 chromosomes at an average depth of 1.2X (*SI Appendix*, Table S3 and Figs. S2-S5), which were used as a platform for subsequent studies.

When compared to the B73 reference genome, Par_N1, Par_N9, and Par_N16 yielded 21,123, 15,554, and 275,990 single nucleotide polymorphisms (SNPs), respectively. To eliminate any potential miscalls caused by postmortem damage, all SNPs corresponding to a possible cytosine (C) to thymine (T), or guanine (G) to adenine (A) transitions were not considered for subsequent analysis (molecular damage filter) (38, 39) (*SI Appendix*, Table S4). All SNPs corresponding to insertions or deletions (INDELs) were also eliminated. Using a previously reported pipeline (9, 30, 31), Par_N1, Par_N9, and Par_N16 yielded 2,886, 1,888, and 121,842 intersected positions with called genotypes (genotype calls) included in the HapMap3 maize diversity panel, most of which were only covered at 1X due to the low amount of endogenous DNA recovered (*SI Appendix*, Table S4). Despite this low coverage depth, the vast majority corresponded to a previously reported HapMap3 allele (98.8% for Par_N1, 98.9% for Par_N9, and 99.1% for Par_N16), suggesting that this dataset provides an accurate paleogenomics representation of maize that can be used to determine its evolutionary trend. It was suggested to us that eliminating specific transitions consistent with molecular damage (C to T and G to A) while keeping others (T to C and A to G) could bias our results in favor of maize alleles. To assess this, we conducted parallel analyses in which we eliminated all transitions (i.e., only transversions were used). However, Par_N16 was the only sample with sufficient genetic information to conduct this second strategy. We obtained 64,118 intersected SNPs involving transversions between this sample and the HapMap3 panel.

### Relationship between ancient maize, extant landraces, and Balsas teosinte

To better understand the origin and domestication of South American maize, we explored the evolutionary relationship between Paredones specimens, teosintes *parviglumis* and *mexicana*, and extant maize landraces. We inferred a bootstrapped maximum-likelihood (ML) tree topology through patterns of population divergence applied to genome-wide polymorphisms. Intersected positions among the three ancient Paredones samples were scarce (*SI Appendix*, Fig. S6); therefore, topologies were constructed individually for each DNA sample, based on the intersection of genotype calls between each of the samples and the maize HapMap3 dataset that includes B73 as a reference genome (including major and minor frequency alleles), 22 maize landraces (including several originating in Mexico), 15 teosinte *parviglumis* inbred lines, two accessions of teosinte *mexicana*, and a single accession of *Tripsacum dactyloides* acting as the outgroup (*SI Appendix*, Table S5). In the case of Par_N16, the resulting tree shows all maize landraces and teosinte accessions separated into two distinct groups, all derived from *Tripsacum* as previously reported (4, 40). Par_N16 is in a clade that includes extant maize landraces, and this is for all 10,000 bootstrap replicates tested. Par_N16 is not basal in its clade but fits robustly with *Chullpi* (AYA 32) – the only extant Peruvian landrace included in the reference panel – in a derived position, closely clustering with South American landraces such as *Cravo Riogranense* (RGSVII) and *Araguito* (VEN 568). These relationships indicate that the ancient samples are monophyletic with modern maize, supporting a single domestication event, and that they are most closely related to modern samples from the same region, strongly suggesting an ancestral relationship between them and modern South American germplasm (Fig. 2 and *SI Appendix*, Fig. S7). In the case of Par_N1 and Par_N9, and although genotype calls intersected with HapMap3 were scarce (2,886 and 1,888, respectively), the resulting topology is equivalent, with both samples clustering at the same position as Par_N16 (*SI Appendix*, Figs. S8 and S9). In general, ancient samples tend to have long branches in phylogenies, which can be explained by isolation by time. On the other hand, the fact that 3 independent samples present the same position in the phylogeny indicates that molecular damage, which is random, is not driving their phylogenetic signal. The parallel analysis in which we used only transversions showed the same topology, where Par_N16 groups with the South American landraces within the maize monophyletic clade (*SI Appendix*, Fig. S10). This shows that the phylogenetic position of the Paredones ancient samples is not biased by the molecular damage filter. Thus, based on genome-wide relatedness, Paredones maize clusters with domesticated Andean landraces.

**Figure 2.**
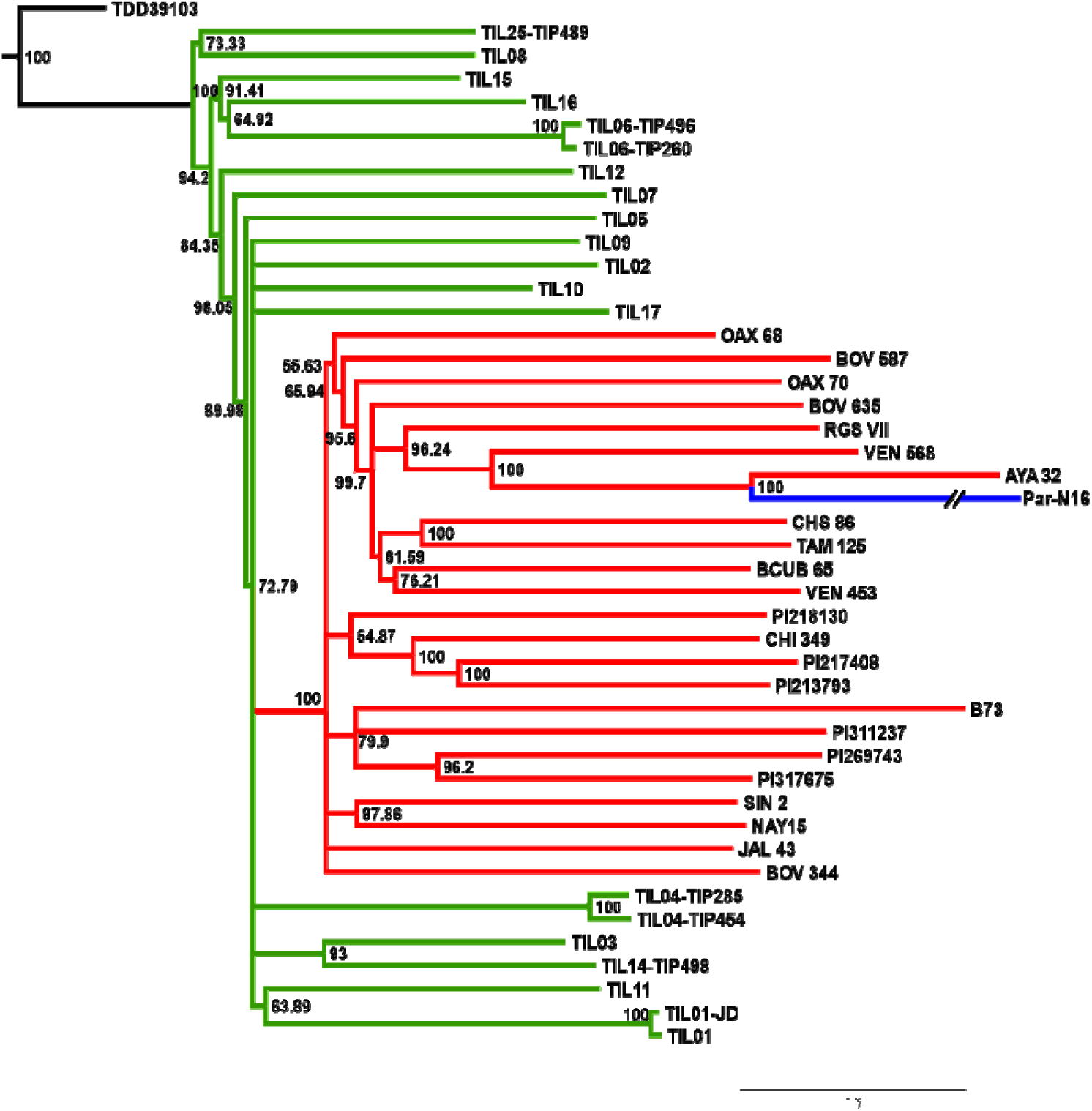
Advanced domestication of ancient Peruvian maize. Evolutionary relationships between ancient Par_N16 maize and its wild and cultivated relatives. ML tree from an alignment of 121,842 genome-wide genotype calls covering non-repetitive regions of the reference maize genome. The teosinte group is highlighted in green, the maize landrace group in red, and the ancient maize sample from Paredones in blue. The teosinte and landrace accessions follow previously reported nomenclatures. The Par_N16 branch was cut for format reasons; a tree with the complete branch can be seen in Fig. S7 (*SI Appendix*).

### *Tests of gene flow from* mexicana

To investigate the genetic relationship of ancient Paredones maize with teosinte *mexicana*, we estimated D-statistics in the form D(*parviglumis, mexicana*, TEST, *Tripsacum*) that test the hypothesis of Incomplete Lineage Sorting (ILS) due to persistence of polymorphisms across different divergence events, against an imbalanced gene flow over derived alleles from *parviglumis* to TEST, and *mexicana* to TEST (*SI Appendix*, Fig. S11). We used highland *Palomero Toluqueño* (PT2233) as a positive control, in the form D(*parviglumis, mexicana*, PT2233, *Tripsacum*), and lowland *Reventador* (BKN022) as a negative control, in the form D(*parviglumis, mexicana*, BKN022, *Tripsacum*). The results of multiple D-statistic distributions show that the positive control PT2233, with D>0, deviated from the balanced gene flow towards *mexicana*. Meanwhile, BKN022 remains in ILS balance, with D around 0. The D-statistics distribution of Par_N16 D(*parviglumis, mexicana*, ParN16, *Tripsacum*) is statistically similar to the distribution of *Reventador* (two-sample Kolmogorov-Smirnov, *p*=0.5814) and significantly different from the distribution of *Palomero Toluqueño* (two-sample Kolmogorov-Smirnov, *p*<0.0001) (*SI Appendix*, Fig. S12 and Table S6). The standard deviation of all 1000 bootstraps is in all cases narrow (SD<0.001), suggesting that the D values are consistent across the genomes. These results agree with a D statistics analysis in which only transversions were used (*SI Appendix*, Fig. S13 and Table S7), showing that the absence of significant gene flow between Par_N16 and *mexicana* is not biased by the molecular damage filter.

To further confirm the absence of *mexicana* introgression in Par_N16 we contrasted the gene flow between *mexicana* and *parviglumis* against the gene flow between *mexicana* and the test sample with D-statistics in the form D(TEST, *parviglumis, mexicana, Tripsacum*) (*SI Appendix*, Fig. S14). In this case, D<0 is an indication of a higher gene flow between *mexicana* and TEST than between *mexicana* and *parviglumis*. As expected, the highland control in the form D(PT2233, *parviglumis, mexicana, Tripsacum*) resulted in D<0, showing significantly higher gene flow with *mexicana* than the one observed between *mexicana* and *parviglumis*. Both the lowlands negative control BKN022 and Par_N16 resulted in D>0, indicating low levels of gene flow between either BKN022 or Par_N16 and *mexicana* (Fig. 3 and *SI Appendix*, Table S8). In both cases, the narrow standard deviation of 1000 bootstrap replications (SD<0.001) suggests that these D values are consistent across the genomes. The result of D>0 for the ancient Paredones maize and the fact that Par_N16 has a significantly higher D value than both lowland and highland controls (two-sample Kolmogorov-Smirnov, *p*<0.0001), was also replicated in the parallel test conducted with transversions only (*SI Appendix*, Fig. S15 and Table S9), showing again that the lower degree of gene flow between *mexicana* and Par_N16 is not biased by the molecular damage filter. These results show that the lineage that gave rise to Paredones maize left Mesoamerica without introgressions from teosinte *mexicana*.

**Figure 3.**
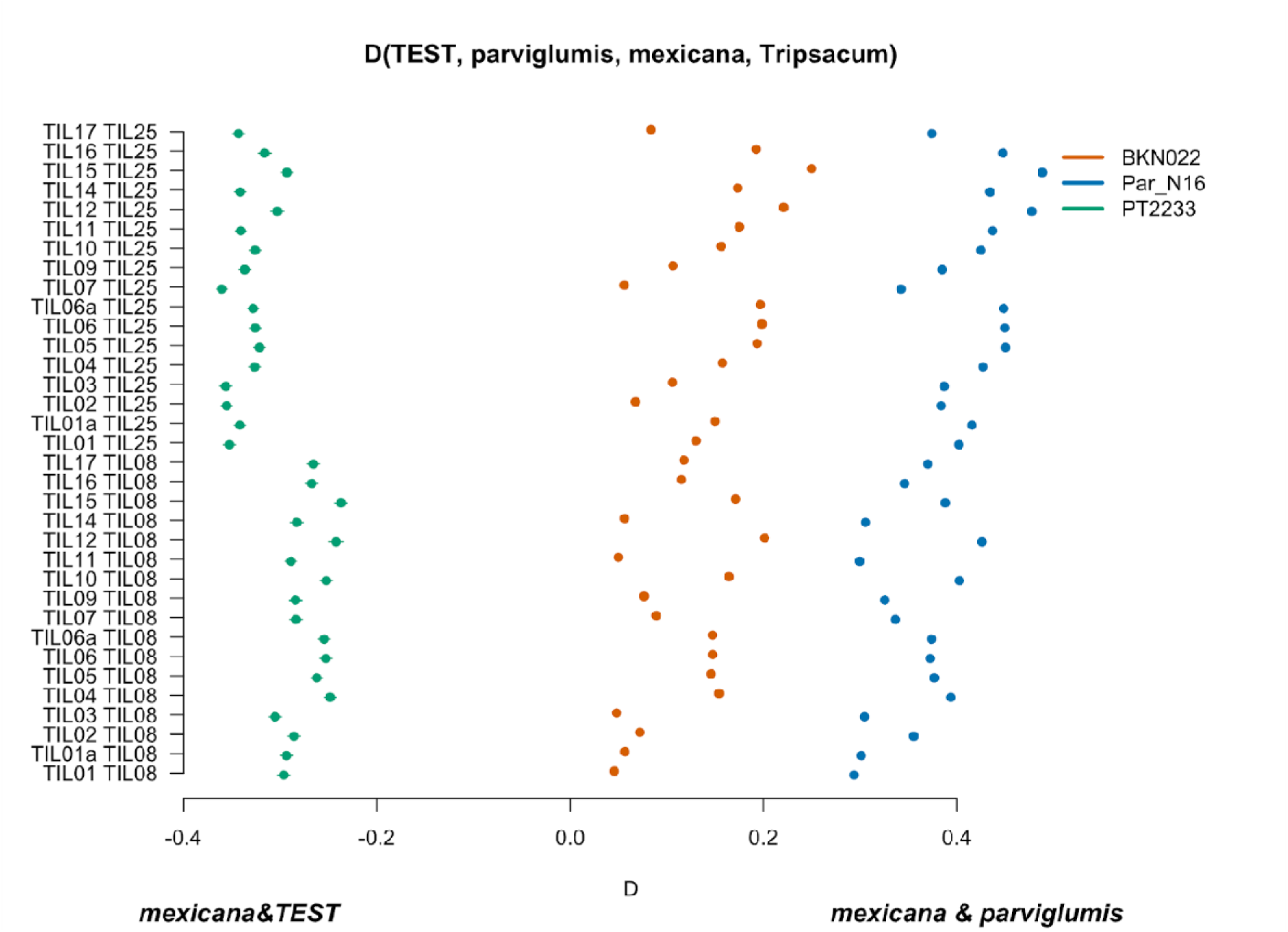
Characterization of *mexicana* gene flow with Par_N16 and Mesoamerican landraces. Genetic comparison of Par_N16 to teosinte *parviglumis* and *mexicana* accessions. D-statistics were calculated in the form D(TEST, *parviglumis, mexicana*, outgroup) (*SI Appendix*, Fig. S14) by comparing 121,842 variant sites shared between Par_N16, *Palomero toluqueño* (PT2233), or *Reventador* (BKN022), and the corresponding SNP variants from teosinte *parviglumis* (TIL01-TIL07, TIL09-17) and two teosinte *mexicana* accessions (TIL25, TIL08). The graph shows the total number of pairwise comparisons (n=34) (*SI Appendix*, Table S8) yielding a negative D for *mexicana* introgression over a test sample or positive D for a higher introgression of *mexicana* and *parviglumis*. Lines in each dot reflect the standard deviation calculated from 100 jackknife replicates.

### Specific adaptation to lowlands in Mesoamerica and South America

Our phylogenetic analysis indicates that the lineage leading to Paredones maize left Mesoamerica already domesticated. However, it does not provide an assessment of how much adaptive variation derives from that acquired in Mesoamerica and how much occurred after moving to South America. To assess this, we identified in Par_N16 all covered SNPs with alleles previously reported to be adaptive to highlands and lowlands, specifically in Mesoamerica or South America (19). From the 668 Mesoamerican and 390 South American previously reported adaptive SNPs, 32 and 20 were covered in Par_N16, respectively. Although at low proportions, the adaptive SNPs in Par_N16 are not significantly underrepresented (*p*=0.8009 and *p*=0.2962, for Mesoamerica and South America, respectively), relative to coverage expectation of non-adaptive SNPs obtained from the same study (19) (see Material and Methods, and *SI Appendix*, Figs. S16). Also, the difference in the proportion of adaptive SNPs covered in Par_N16 and corresponding to Mesoamerica and South America is not statistically significant (32/668 vs. 20/390, Fisher exact test, *p*=0.8832), suggesting an equivalent power of detection within both SNP sets. The similarity between Par_N16 and highland or lowland populations in Mesoamerica and South America can be quantified as the average allelic similarity between the ancient sample and the respective modern populations at a given set of SNPs. Comparison of similarity estimated from SNPs previously reported as adaptive to similarities estimated from multiple random samples of background SNPs allows a quantification of the deviations from genome-wide similarity expectations (see Material and Methods, Fig. 4, and *SI Appendix*, Tables S10 and S11). The mean genome-wide relatedness of Par_N16 is 0.785 for highland and 0.800 for lowland populations in Mesoamerica, and 0.831 for highland and 0.812 for lowland populations in South America (Fig. 4). Thus, at the genome-wide level, Par_N16 is genetically more similar to South American landraces, particularly from the highlands, than to Mesoamerican populations. At adaptive loci, Par_N16 has lower proportions of shared alleles to highlands than to lowlands for both Mesoamerica and South America (Fig. 4). In adaptive loci, for Mesoamerica, Par_N16 has an average similarity with highland genotypes of 0.4995 and 0.7436 with lowland individuals (*SI Appendix*, Datasets S1 and S2), while for South America the similarity with highland genotypes was 0.4918, compared to 0.705 for the lowlands (*SI Appendix*, Datasets S3 and S4). Moreover, Par_N16 is significantly less similar in adaptive SNPs with the highland populations from both regions relative to genome-wide expectations (*p*<0.0001 in both cases); however, its adaptive similarity with lowland populations was significantly reduced for South America (*p*=0.0386) but not for Mesoamerica (*p*=0.1116). Nevertheless, haplotype sharing for both lowland populations is not far outside of genome-wide expectations, in contrast with highland populations (Fig. 4). Thus, although Par_N16 is still more adapted to lowland Mesoamerica, it was in the process of adapting to lowland South America.

**Figure 4.**
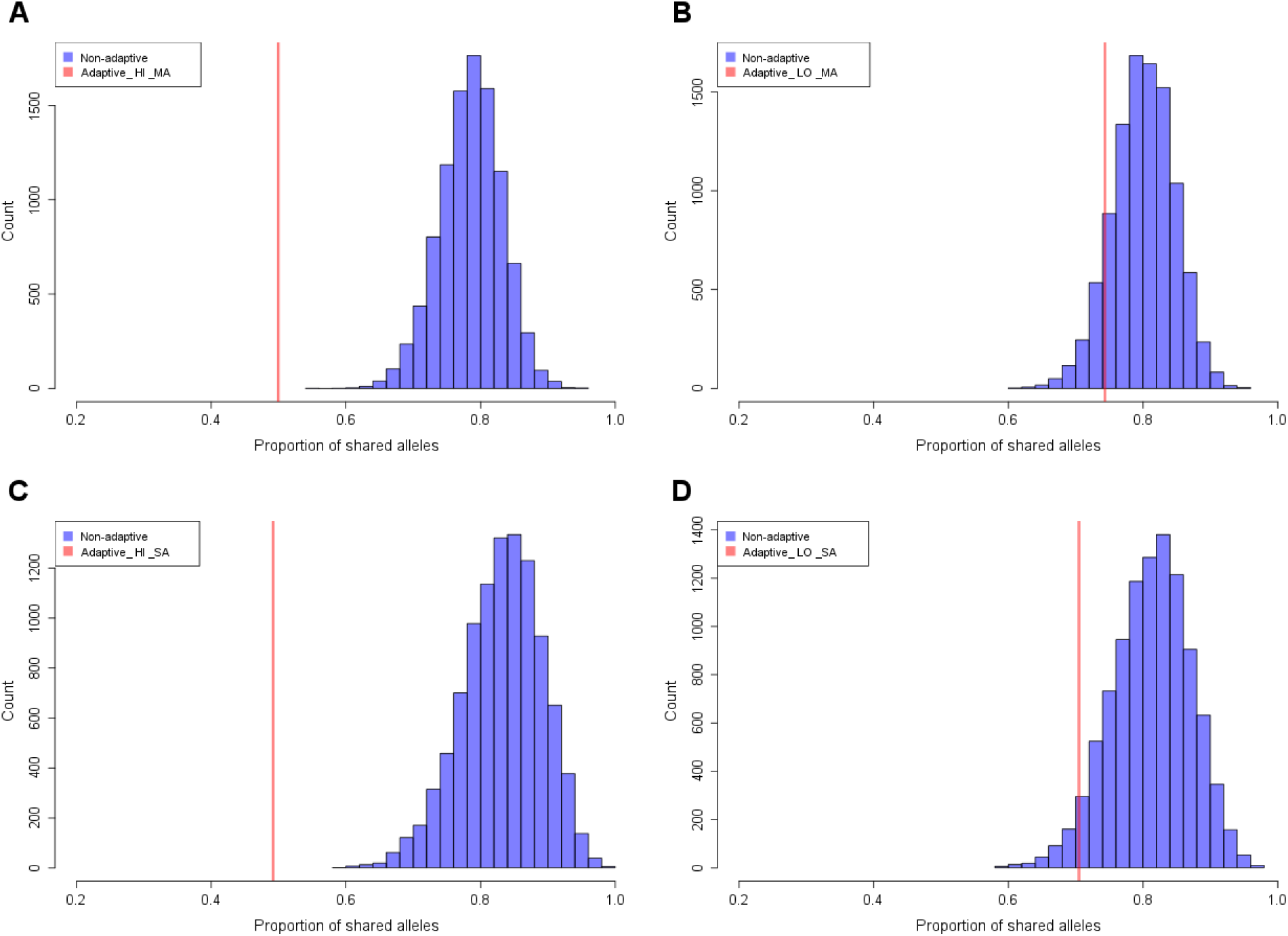
Allelic similarity between Par_N16 and landraces from Mesoamerica (MA) and South America (SA). Comparisons involved genome-wide non-adaptive SNPs (blue distributions) and SNPs with significant *F*_*ST*_ implicated as adaptive (red lines) at intersected sites between Par_N16 and the reference dataset (19) (see Material and Methods). **A**, the expected average in genome-wide allelic similarity between Par_N16 and highland MA landraces in non-adaptive SNPs is 0.785; the corresponding allelic similarity in adaptive SNPs is 0.4995. **B**, the expected average in genome-wide allelic similarity between Par_N16 and lowland MA landraces in non-adaptive SNPs is 0.8; the corresponding allelic similarity in adaptive SNPs is 0.7436. **C**, the expected average in genome-wide allelic similarity between Par_N16 and highland SA landraces in non-adaptive SNPs is 0.831; the corresponding allelic similarity in adaptive SNPs is 0.4918. **D**, the expected average in genome-wide allelic similarity between Par_N16 and lowland SA landraces in non-adaptive SNPs is 0.812; the corresponding allelic similarity in adaptive SNPs is 0.705 (Datasets S1-S4).

## Discussion

Paredones ancient maize represents the earliest known macro-specimens of maize and was found in Peru 3,800 km away from the center of origin. Paredones samples are morphologically similar to extant maize while the earliest maize from Mexico still retained shared morphology and haplotypic diversity with wild populations. Therefore, the recovery of genomic sequences of these early South American populations brings a unique opportunity to reconstruct the adaptation and dispersal processes of maize. Considering the antiquity and location of these samples, the genomic information we obtained is valuable. All three samples analyzed here are located within the monophyletic clade of maize, indicating that the ultimate origin of ancient Paredones maize is not different from all Mexican landraces examined to date and supporting a single domestication event. The ancient samples were grouped in all cases within a subclade of South American landraces, particularly with the Peruvian landrace Chullpi, suggesting that ancient Paredones maize was already domesticated by 6775 to 5324 calibrated BP and at least partially ancestral to extant South American landraces. Previous genomic analyses from ancient and modern maize from South America were interpreted as evidence of stratified domestication, in which one of several partially domesticated lineages arrived early (at least 7,000 BP) to South America and locally evolved all domestication traits (21, 30). An ancient sample located in an ancestral or sister position to the Mesoamerican maize clade would provide evidence to support this model. The phylogenetic position of Paredones samples does not show this pattern (Fig. 2) and does not support the stratified model as previously proposed (21, 30), but it is compatible with a sequential model of crop evolution in which domestication is the first stage, followed by an increase in frequency of desirable alleles (stage 2), and the formation of cultivated populations adapted to new environments and local preferences (stage 3) (41).

In Mesoamerica, maize adaptation to highlands was marked by the introgression of alleles from the highlands teosinte *mexicana*, and maize lineages adapted to Mesoamerican highlands carry this gene flow signal (11, 13). Previous research suggested that South American highland maize was independently adapted from local lowland germplasm rather than relying on the same allelic diversity that underlies highland adaptation in Mexico (19). Our analyses show that there was no significant gene flow between Par_N16 and *mexicana*. If any, it was significantly lower than the gene flow between *mexicana* and lowland landraces from Mexico such as *Reventador*, and also significantly lower than the gene flow between *parviglumis* and *mexicana*. This result suggests that the early Paredones maize populations diverged from Mesoamerica without gene flow from *mexicana* or any highlands maize in Mexico, consistent with the idea that *mexicana* introgression into maize populations occurred more recently (1000 generations ago) (15). While modern highland South American germplasm shows evidence of *mexicana* introgression (10), Paredones material does not contain *mexicana* allelic diversity and it is possible that the earliest germplasm that was grown in the Andes did not contain it either. This raises the possibility that there is novel highland adaptive diversity harbored by South American landraces, however, more ancient and modern samples, especially from highland Andean locations, are needed to test this. In addition, *mexicana* introgression is pervasive across domesticated maize (10, 15); therefore, Paredones ancient samples might be useful as minimal or no-introgression controls in future studies assessing *mexicana*-maize gene flow.

At the genome-wide level, Par_N16 is more similar to the lowland than to the highland Mesoamerican population (Fig. 4). Meanwhile, in South America, the genome-wide similarity is higher than in Mesoamerica but especially in the highlands, which is interesting because Paredones is a lowland site. One possible explanation is that Paredones is likely ancestral to both lowland and highland populations (with the latter derived from local lowland landraces), but that subsequent gene flow from Mesoamerica (10, 19) had a greater impact on lowland populations, erasing part of this ancestry. Understanding the process of highland adaptation will require additional sampling in both the highlands and the lowlands over time.

Allele similarity at SNPs that showed significant *F*_*ST*_ values between the highlands and the lowlands in Mesoamerica and South America (19), clearly shows that the Paredones sample has far less similarity with the highland populations (*p*<0.0001), as is consistent with their lowland provenance; but surprisingly, they share a higher proportion of adaptive SNPs with lowland Mesoamerican populations than with lowland South American ones, for which similarity was significantly reduced (*p*=0.0386). These results seem to suggest that ancient Paredones samples were better adapted to their ancestral Mexican lowlands than to their new environment in lowland South America. However, Par_N16 still shares some similarities to lowland South America in adaptive SNPs (unlike highland), evidencing some level of adaptation to the new environment. In addition, the deficiency of adaptive alleles in this region can be explained if an important part of the current adaptive alleles were to arrive or increase in frequency in the ancestral or later populations in South America at the time of Paredones. Under this perspective, the date of Par_N16 (5583-5324 calibrated BP) suggests that a substantial amount of improvement occurred rapidly and specifically in South American lowlands. However, more sampling over time, and contextualized by archaeological inference into the role of maize in society, is needed to understand how rapidly maize adapts. On the other hand, highland-adaptive alleles are expected to be deleterious in lowlands (19), which could explain their rarity in a lowland sample. All of this places the geographic origin of Paredones allelic diversity in Mesoamerica and supports a lowland coastal migration route. This evidence also suggests that Paredones lineage was in stage 3 of the crop evolution model referred to above (41). In the end, adaptations and improvements occurring in both Mesoamerica and South America can explain the rapid evolution responsible for the modern phenotype that Paredones maize presents despite its antiquity.

In all, our results suggest that, unlike in the highlands (8, 9), domestication occurred in lowlands Mexico before Paredones lineage arrived in South America following a coastal Pacific corridor of cultural and physical goods from Mesoamerica to Peru. Under this scenario, domestication and improvements in Mesoamerican lowlands, migration from there to Peru, and further processes of local adaptation must have occurred throughout ∼2500 years, assuming a teosinte-maize divergence time of 9,000 years (4). During this relatively short period, there was no gene flow between *mexicana* and this maize lineage, but there were processes of specific adaptation to South American lowlands, which required expert management in the face of new environmental and socio-economic pressures. This fits well within the unique and advanced developments of the Andean societies that occurred during a period of rapid cultural transformation between 7500 and 6500 calibrated BP, involving permanent agro-maritime villages along the Pacific shoreline, farming communities in coastal and highland valleys, camelid husbandry, monumental architecture and public ritual, elaborate art and iconography, and craft production (17, 42–44). These and other developments occurred earlier and were more widespread than elsewhere in the hemisphere, resulting in the establishment of new social organizations, increased plant cultivation, and intensive landscape modification. Collectively, these mid-Holocene transformations represent a package of social and cultural traits unforeseen at any other region of the Americas.

## Materials and Methods

Detailed descriptions of samples and methods are provided in the *SI Appendix*.

### Samples

Paredones ancient specimens and date determination were previously reported (27, 28) (SI Appendix). Sample processing and DNA extraction were performed following all necessary procedures to avoid human-related or cross-sample contamination in a clean Laboratory optimized for paleogenomics, as previously described (8).

### Sequencing of ancient samples

Two double-index DNA Illumina libraries for each sample were built at Max Planck Institute Tübingen, using established methodologies for ancient DNA (10, 45–47). From a second Par_N16 library, DNA fragments of 150 to 205 bp in length were selected using the Pippin Prep DNA size selection system (Sage Science) with 2% agarose gel cassettes. Subsequent shotgun sequencing of Illumina libraries was conducted with Nextseq at Unidad de Genómica Avanzada, Laboratorio Nacional de Genómica para la Biodiversidad, Cinvestav Irapuato.

### Read processing, mapping, and genotyping

Custom double-index sequences of 8 nucleotides were used to tag the libraries described above. All libraries were filtered to remove adaptors and low-quality reads using Cutadapt (48). Rescaling of Phred quality scores to account for molecular damage was implemented using mapDamage2.2 (49) and keeping reads longer than 30 bp with a quality above 10 Phred score. Filtered reads were mapped against *Z. mays* B73 RefGen_v3 (50) using the Burrows-Wheeler analysis (BWA) MEM algorithm with default conditions (51).

### Metagenomic analysis and post-mortem damage

Cytosine deamination rates and fragmentation patterns were estimated using mapDamage2.2 (49). All sites behaving as molecular damage (CG→TA) were excluded. Heterozygous sites with one damage variant were transformed into homozygous sites for the non-damaged variant. A metagenomic filter was applied to discard reads that aligned to sequences in the GenBank National Center for Biotechnology Information database of all bacterial and fungal genomes using default mapping-quality parameters of BWA (44). Parallel analyses were conducted using only transversions to assess potential bias introduced by our molecular damage filter.

### Evolutionary analysis and genotype comparisons

Patterns of divergence were analyzed by generating ML trees using Treemix (52) and the intersection of genotype calls between the ancient specimens and 44 selected individuals of the publicly available database HapMap3 without imputation (53). For each tree, 10,000 bootstrap pseudo-replicates were generated with a parallelized version of a public script (https://github.com/mgharvey/misc_python/blob/master/bin/TreeMix/treemix_tree_with_bootstraps.py), which uses the *sumtree* function in *DendroPy* (54) to obtain a consensus ML bootstrapped tree.

### Introgression analysis

Quantification of *mexicana* introgression was performed by D-statistics in form D(P1, P2, P3, O) calculated from an ABBA(*x*) BABA(*y*) scheme *D=(x-y)/(x+y)*; being *x* the total amount of haplotypes shared between P2 and P3, and *y* the total amount of haplotypes shared between P1 and P3. We used CalcD from the evobiR tools package (55) performing 100 Jackknife replicates. We used all genotype calls intersected between test samples and HapMap3 (53). We used a Wilcoxon nonparametric test for testing differences between positive and negative values. Two-sample Kolmogorov-Smirnov (K-S) tests were conducted using ks.test R package version 1.2.

### Coverage at adaptive sites

To assess the significance of the coverage at adaptive alleles, we generated 10,000 lists of 668 for Mesoamerica or 390 for South America SNPs each, which were randomly sampled from 63,271 non-adaptive SNPs intersected between Par_N16 and the public dataset (19). We recorded the number of SNPs from each of the 668 or 390 lists that were covered in Par_N16, obtaining the respective null coverage distributions that reflect the coverage expectation of sampling 668 or 390 SNPs in Par_N16. The probability of underrepresentation of covered SNPs or adaptive alleles in Par_N16 is then the proportion of the null distribution (n=10,000) that showed the same value or less than that observed in Par_N16 for the 668 or 390 adaptive SNPs.

### Adaptation to Mesoamerican and South American lands

The reference (published) data consisted of 668 SNPs specific to Mesoamerica and 390 SNPs specific to South America with significant *F*_*ST*_ values between highland and lowland populations and therefore considered to be adaptive, as well as 647,821 non-adaptive SNPs (without significant *F*_*ST*_ values) from the analyzed panel (19). Covered SNPs in Par_N16 were identified by the intersection with the above-mentioned adaptive and non-adaptive SNPs. Allelic similarity in adaptive SNPs was obtained by calculating the mean genetic distances between Par_N16 and each of the four test populations (highland Mesoamerica, highland South America, lowland Mesoamerica and lowland South America) at the Intersected sites. To assess significance, we generated null distributions of genome-wide similarity expectations for each of the four test populations. We generated 10,000 random samples from the 63,271 non-adaptive SNPs covered in Par_N16, obtaining the mean genetic distance for each sample. Each random sample consisted in the same number of SNPs as the number of adaptive SNPs covered in Par_N16 (32 for Mesoamerica and 20 for South America). The statistical significance of the reduction in adaptive similarity relative to genome-wide similarity is then the proportion of the null distribution (n=10,000) that showed the same or less similarity than that observed in Par_N16 for the 32 or 20 covered adaptive SNPs.

## Acknowledgments

We would like to thank Hilda Ramos-Aboites and Christian Martinez-Guerrero for technical support, as well as Peruvian archeologists and the Ministry of Culture in Lima, Peru, for granting permission to carry out the archeological research and analysis. We thank Hernán A. Burbano for advice on molecular biology protocols and for hosting M.V-E. during his visit to Germany. The GBS data from South American landraces (19) used in the adaptation analysis were kindly provided by Jeffrey Ross-Ibarra. M.V-E., G.G.H-R., E.G-O. and I.L-V were recipients of a graduate scholarship from Consejo Nacional de Ciencia y Tecnología (CONACyT). This research was supported by a CONACyT grant to J.P.V-C. (CB256826), the INAH-Cinvestav Tehuacan initiative, and Cinvestav internal funds assigned to R.M.

## Data deposition

Sequence data generated for this study can be accessed by direct contact to corresponding authors and will be deposited in the European Nucleotide Archive upon final publication.

