## Supplementary Information for "Domestication and lowland adaptation of coastal preceramic maize from Paredones, Peru"


**This PDF file includes:**

Supplementary text
Figures S1 to S17
Tables S1 to S11
Legends for Datasets S1 to S4
SI References

**Other supplementary materials for this manuscript include the following:**

Datasets S1 to S4 in a single Excel file

### Supplementary Text

#### Materials and Methods

#### Radiocarbon Dates on Maize Macro-Remains: Chronology and Stratigraphy.

**1**. The chrono-stratigraphic data for the maize remains were published previously in Grobman *et al*. in 2012 (1). These data come primarily from archeological excavations at the Huaca Prieta and Paredones sites, both multicomponent Preceramic-aged localities situated on a remnant Pleistocene terrace overlooking the Pacific Ocean on the north coast of Peru. Huaca Prieta is a large artificial earthen and stone mound approximately 165 m long, 85 m wide, and 30 m high. Paredones is an artificial earthen mound and measures approximately 40 long, 23 m wide, and 6 m high. The macro-maize remains from both sites were excavated in deeply stratified, intact (i.e., undisturbed) cultural floors. Stratigraphic Unit 22 at Paredones is the archeological component with the largest and most diversified amount of maize remains, with the oldest $C^{14}$ dated cobs derived from the base of this unit (see Fig. 1), in a single, discrete and intact floor of ~2 cm in thickness and at 5.5 m in depth from the present-day surface (2).

AMS radiocarbon ages on the maize remains were obtained on a burned shank and on unburned and burned cobs by laboratories at the University of Arizona, Beta Analytic Inc., and the Woods Hole Oceanographic Institute. All dates were calibrated using shcal04 (3). Furthermore, the dated remains at both sites are chrono-stratigraphically bracketed by and in agreement with more than 160 AMS and OSL dates from the mound and off-mound contexts (2). This allowed the project to cross-reference and control multiple, wide, and deep stratigraphic units within and across excavation units at sites, and to document any taphonomic and other potential disturbances that might have affected the integrity of the context and absolute age of the remains. As discussed below, no taphonomic or other disturbing cultural or geological features were observed in any excavation units that would have altered the integrity and intactness of strata containing the maize remains. Except for five anomalous assays, the vast majority of the dates conform to their expected chronological order and stratigraphic position; and agree with the few radiocarbon measures obtained by Bird (4) in the 1950s.

All 145+ $C^{14}$ dates were measured on single chunks of wood charcoal, animal bone, corn remains, and cotton textiles recovered from intact hearths, shallow food pits (2-3cm thick), and human burials embedded in floors. All wood charcoal dates were derived from short-lived bushes and small short-lived trees. No radiocarbon samples were taken from fills, middens, and marine shells. Given the different organic materials dated by four different laboratories over a period of six decades (including Bird's dates by Libby's laboratory in the 1950s), all dates generally agree and overlap chronologically and stratigraphically at the 2 sigma or 98.2% age range calibrated for floor sequences in stratigraphic units.

The most complete stratigraphic sequence on dated macro-remains is a series of AMS assays obtained from hearths and shallow features in intact floors from Unit 22 at Paredones. A charred cob fragment from Floor 6 at a depth of 1.2 m was dated at 4821–4527 cal BP (AA86934). Dates on single chunks of wood charcoal from hearths embedded in Floors 10, 15, and 16 at depths of 3.8m, 4.7m, and 4.9m, respectively, were processed at 5435–5044 cal BP (Beta263320), 5585–5325 cal BP (Beta263321), and 5711–5335 cal BP (AA86947). Recently, a charred cob fragment from Floor 11 was assayed at 5404-4937 cal BP (D-AMS 044318), which agrees with its chrono-stratigraphic position and other AMS dates in the site. An uncharred cob fragment directly below Floor 10, at a depth of ~4.0m, was dated at 133–34 cal BP (Beta263988). This date is anomalous and incorrect. An articulated husk and charred shank fragment from Floor 18, at a depth of 5.5m in Unit 22, yielded a date of 6775–6504 cal BP (OS86020). Underlying this date at a depth of 5.8m is an assay of 6640–6319 cal BP (AA83260) on wood charcoal from Floor 24, Unit 22. Four assays were processed on an uncharred cob segment from Floor 18. These were anomalous dates that were younger, including one post-bomb date than the charred shank fragment that dated at 6775–6504 cal BP on the same floor. Dates on the husk and charred shank fragment from Floor 18 indicate that the harder, more durable husk and charred shank yielded a reliable $C^{14}$ measure that completely agrees with its stratigraphic position below the assays in Floors 10, 15, and 16 and above the one in Floor 24. It also indicates that the assays on unburned, soft fragments of cob are too young and thus unreliable. In sum, the dates on wood charcoal and charred shank and cobs from Unit 22 are in complete stratigraphic agreement, showing that the four anomalous assays on the one unburned cob in Floor 18 are in error.

In 2019, Dillehay and geologists Steven Goodbred and Elizabeth Chamberlain carried out excavations in a Preceramic domestic site (S-18) located ~3.2 km north of Huaca Prieta. Preceramic corn remains were encountered consistently in the upper to lower intact cultural layers of the site. As with the Paredones and Huaca Prieta sites, the lower strata contained both charred and uncharred cobs of the smaller and earliest type of identified corn species, Proto Confite Morocho (1) (Grobman, personal communication, 2019). The middle to upper strata yielded the known later and slightly larger Preceramic varieties of Confite Chavinese and Proto Alazan. An OSL date from a discrete and intact lower layer containing a hearth with two unburned cob fragments of the Proto Confite Morocho variety assayed at ~7000 +/- 630 years ago or 5610-4350 BCE (5). Wood charcoal from the hearth processed at 7162-6914 +/- 30 cal BP (AA75398).

Other dates on burned cob fragments from other localities in and around Paredones and Huaca Prieta also were in chrono-stratigraphic order and dated 4149–3839 cal BP (Beta278050), 3956–3704 cal BP (AA86941), 3982– 3728 cal BP (AA86931), and 4235–3928 cal BP (AA86946).

In South America, maize micro remains (e.g. starch grains, pollen, phytoliths) have been dated ~7500–7000 cal BP (6–8) at sites in southwest coastal Ecuador, located ~450km north of Huaca Prieta and Paredones, and in other localities across the continent dated ~6500 cal BP and later (9).

**2**. In specific regard to the four anomalous dates from Floor 18 in Unit 22, there is no taphonomic or stratigraphic evidence (e.g., pits, animal burrows, tree roots, postholes, truncated strata) to indicate intrusiveness of younger materials or post-occupation disturbance in any of the excavated contexts at Huaca Prieta and Paredones (see Fig. 1 for the discrete, undisturbed floors and fills and the location of AMS assays on macro-corn remains and wood charcoal from Unit 22 at Paredones). Moreover, all strike and dip measurements taken on piece-plotted artifacts and features embedded in floors show no upward or downward tilting indicative of post-depositional disturbance, that is, all stone tools, bone, shell, and plants remains and other ecological and artifactual debris laid flat in or on floors, indicating undisturbed and unimpeachable contexts.

The anomalous dates on uncharred cob remains from Floor 18 range from ~600-700 cal BP to recent post-bomb years. With regard to their discrepancy with the 6775–6504 cal BP on a fragment of husk and charred shank, a younger carbon-bearing element must have penetrated the attached unburned cob fragment and contaminated it.

Unburned cobs with younger dates may have been contaminated by mold, fungus, heavy salt saturation from the local seashore environment, or some other element that might have affected them. We performed SEM analyses of the microscopic cellular structure of unburned cobs, husk, and shanks and of wood charcoal. Fungal activity was identified only in the cellular structure of unburned cobs (personal communication with Duccio Bonavia, 2011, Faculttad de Biologia, Universidad Cayetano Heredia). No fungal activity was observed in the cells of wood charcoal and of the husk and charred shank. With regard to possible contamination effects from fungi, Darden Hood (Beta Analytic Inc., Miami, FL) in a personal communication to Dillehay in 2011 proposed reasonable causes to consider: "...uncharred corn acts like a sponge and its integrity is too weak to withstand the pretreatments prior to removing organic contaminants. Thus, the radiocarbon (RC) pretreatments are dissolving the sample just as fast as contamination, resulting simply in a reduction in sample size rather than de-contamination" and "...if the corn was being preferentially removed with the alkali, thereby increasing the concentration of the fungus, the date would come out younger with a higher fungus to corn ratio. As you go deeper [stratigraphically], the corn is more weathered, and more susceptible to removal with the alkali... whereas the fungus is fresh-and-resistant."

Certain conclusions can be drawn from the anomalous dates on the unburned cob. There is no evidence of post-depositional disturbance at any of the excavated sites, including Unit 22. Thus, there is no possibility that the younger dated samples are intrusive. All of the dates considered reliable are entirely coherent within the radiocarbon-dated stratigraphic sequences at Huaca Prieta and Paredones and with directly associated wood charcoal dates from the same feature and floor contexts yielding the maize remains. The most reliable dates are on hard husks and charred shanks and cobs, which have a more rigid, impenetrable cellular structure. This suggests the probability that the other, more porous tissues of soft, uncharred cobs can absorb or allow the growth of some contaminating substance that does not affect the harder tissues of the charred tissue and husks of the maize. Regardless of our suspicions of what might have contaminated a few unburned maize cobs, the archeological contexts of the deeply buried and culturally intact stratigraphic deposits, where all maize remains and acceptable maize and wood dates were recovered, take precedent over the younger, anomalous cob dates.

**3**. The three samples analyzed here consist of well-preserved macro-specimens of maize that were morphologically reminiscent of extant structured maize cobs. Par_N1 was dated directly to 6775-6504 calibrated BP by dating the husk and charred shank; Par_N1 dating number is

OS86020, as referenced in (3). Par_N9 was dated stratigraphically to 5800-5400 calibrated BP, a more accurate and conservative date than the one presented in (3), due to in-depth subsequent dating of its stratigraphic location. Par_N16 was dated indirectly to 5,583-5,324 calibrated BP by association to a piece of wood charcoal found in the same unit U20, stratum (5b). Its dating number is AA86936, as referenced in (3).

#### Extraction and sequencing of Paredones archeological maize samples

All three maize specimens were sampled using forceps and sterile scalpels. DNA extraction was of 14.2 to 15 mg of the inner parenchyma tissues at the ancient DNA facilities of UGA Langebio CINVESTAV.

Isolation of DNA was carried out in clean laboratory facilities at UGA Langebio and Tuebingen University, Germany, with dedicated reagents and equipment that are frequently sterilized, and UV treated. To prevent cross-sample and human-related contamination, we used new disposable plastic material and filtered pipette tips, and personnel wear protective gear such as full bodysuits, masks, and doubled gloves. Work was conducted in laminar flow hoods. Samples were ground with a mortar and pestle. DNA extraction was conducted using a freshly prepared PTB lysis buffer (PTB 2.5mM, DTT 50mM, Proteinase K 0.4mg/ml, 1% SDS, 10 mM Tris, 10 mM EDTA, 5mM NaCl) and purified using QIAgen DNEasy® Plant Mini kit columns following an established protocol (10, 11). All shotgun libraries were constructed from 20 µl of ancient maize DNA following a published protocol tested in maize (10, 12) with modifications as suggested in (13). Libraries were amplified for 10 cycles with unique combinations of two indexing primers (14). The quality of libraries was tested using Qubit 2.0 fluorometer (Thermo Fisher) and a High Sensitivity DNA Assay Chip Kit (Agilent, Waldborn Germany) on a Bioanalyzer 2100 (Agilent Technologies).

Non-UDG double index DNA Illumina libraries for each sample were built at Max Planck Institute Tuebingen, using established methodologies for ancient DNA, for subsequent shotgun sequencing. Illumina libraries were sequenced in three different rounds for a total of 83.88 Gb (Par_N1 35.7 Gb, Par_N9 23.48 Gb, Par_N16 24.7 Gb) using Nextseq at UGA Langebio, Cinvestav.

To increase the endogenous content of the Par_N16 sample a secondary library was sequenced using Nextseq yielding 21.59 of additional Gb. For this library, which originated from a split of the original library already amplified, a final size-selection step was performed using a Pippin-Prep procedure in a 2% agarose cassette (Sage science, Beverly MA) to retain DNA fragments from 150 to 205 bp in size.

#### Read processing, mapping, and genotyping

Double Index sequences of 8 nucleotides were used to tag libraries described above. Only reads with the correct index were used in downstream analysis. All libraries were filtered to remove adaptors and low-quality reads using Cutadapt (V1.13) (15) and keeping reads longer than 30 bp with a quality above 10 Phred score. Repetitive adenines (A) and thymines (T) were invariably removed from read ends. Filtered reads were mapped using the BWA MEM algorithm with default conditions (V0.7.12) (16). *Zea mays* B73 RefGen_v3 (17). Reads with multiple hits were removed using SAMtools map quality filters. As a clonal removal strategy, sequence duplication in reads was filtered with the rmdup function of SAMtools (V1.5) (18), and sequences were locally re-aligned around insertion/deletions (indels) using GATK IndelRealigner (V3.7)(19). Rescaling of phred quality scores to account for molecular damage was implemented by using the --rescale parameter in mapDamage(V2.2) (20). A map quality filter of a minimum value of 10 Phred score was applied to eliminate reads with low certainty assignments. Mapping efficiency of Paredones ancient samples was 0.1409% for Par_N1 and 0.1612% for Par_N9 and 0.728% for Par_N16. SNP and genotype calling were performed as previously described (21, 22). Variation information was extracted and called using the mpileup and bcftools functions of SAMtools (18), generating a VCF file containing genotypes. This entire pipeline has been already optimized for analyzing ancient maize samples (23).

#### Metagenomic analysis and postmortem damage

Cytosine deamination rates and fragmentation patterns were estimated using mapDamage2.2 (20) based on all reads mapping to the B73 reference genome, revealing expected patterns of postmortem damage in the form of C>T substitutions at the 5' termini, and G>A substitutions at the 3' termini. The excess of purines observed near read termini supports fragmentation driven by depurination (Figure S1). All indels and sites behaving as molecular damage (CG->TA) (24, 25) were excluded (molecular damage filter). However, heterozygous sites with one variant compatible with molecular damage were transformed into homozygous sites for the variant without damage pattern. A metagenomic filter was applied to discard reads that aligned to sequences in the GenBank NCBI database of all bacterial and fungal genomes using default mapping quality parameters of BWA (16). Parallel analyses were conducted using only transversions to assess potential bias introduced by the molecular damage filter.

#### Evolutionary analysis and SNP genotype comparisons

Patterns of divergence were analyzed by generating maximum likelihood (ML) trees using Treemix (V1.12)(26) and the intersection of SNPs passing quality filters for the ancient specimens and 44 selected individuals of the publicly available database HapMap3 without imputation (27). The list of selected individuals is presented in Table S5. The topologies were generated with each ancient sample individually or including both samples together. In each case, no less than 10,000 bootstrap pseudo-replicates were generated with a parallelized version of a public script (28), which uses the sumtree function in DendroPy (29) to obtain a consensus ML bootstrapped tree. The same SNP alignments were also used to assign the identity of each ancient SNP genotype to shared or exclusive SNP genotypes of the selected HapMap3 individuals. According to their SNP identity, the ancient genotypes were assigned exclusively to one of six categories: B73 genotypes, maize landraces genotypes, *Zea mays* ssp. *parviglumis*, *Zea mays* ssp. *mexicana*, *Tripsacum dactyloides*, or those not present in the dataset (ancient sample's private SNPs). Tree topologies were generated on the basis of an intersection between maize HapMap3 concatenated for each genotype. The resulting trees were visualized using figtree software https://github.com/rambaut/figtree/.

#### Introgression analysis

Quantification of genetic similarity attributable to shared ancestry was performed by D-statistics D (P1, P2, P3, O) calculated from an ABBA($x$) BABA($y$) scheme $D=(x-y)/(x+y)$; being $x$ the total amount of haplotypes shared between P2 and P3, and $y$ the total amount of haplotypes shared between P1 and P3. We used the CalcD function of the evobiR tools package (30, 31), performing 100 jackknife replicates using the parameters "sig.test=J" and "replicate=100" (see https://github.com/coleoguy/evobir). Only sites covered in Par_N16 were considered in the controls. To test genetic similarity to highland and lowlands populations we used GBS public data of South American landraces (32). To test *mexicana* introgression we used all genotype calls intersected between test samples and HapMap3 (27). Two D forms were tested: (i) D(*parviglumis*, *mexicana*, TEST, outgroup) (Figure S11) that allows a direct estimation of gene flow of derived alleles back from *parviglumis* or *mexicana* into the test sample, and (ii) D(TEST, *parviglumis*, *mexicana*, outgroup) (Figure S14) that measures an excess of gene flow between *mexicana* and test compared to the gene flow observed in *mexicana* and *parviglumis.* In both cases *Tripsacum dactyloides* (TDD39103) was set as an outgroup. We used a Wilcoxon nonparametric test for testing differences between positive and negative values using R package 3.6.2 https://www.rdocumentation.org/packages/stats/versions/3.6.2/topics/wilcox.test. Two sample Kolmogorov-Smirnov (K-S) tests were conducted using ks.test R package version 1.2. https://www.rdocumentation.org/packages/dgof/versions/1.2/topics/ks.test.

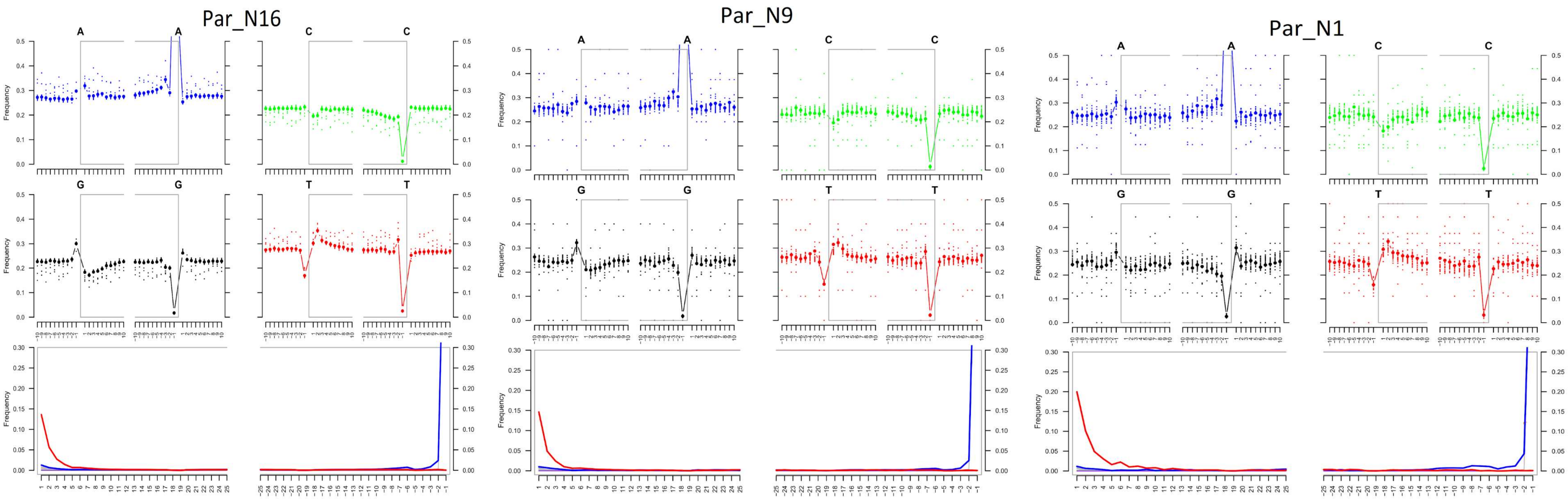


**Fig. S1. Post-mortem DNA damage and fragmentation patterns of ancient maize samples.** DNA composition around read-termini (top four plots), and DNA mis-incorporation errors relative to the 5' and 3' read (bottom plot); the two distributions for post-mortem damage signatures (C to T and G to A) are shown in red and blue respectively, while other types of substitutions are shown in gray.

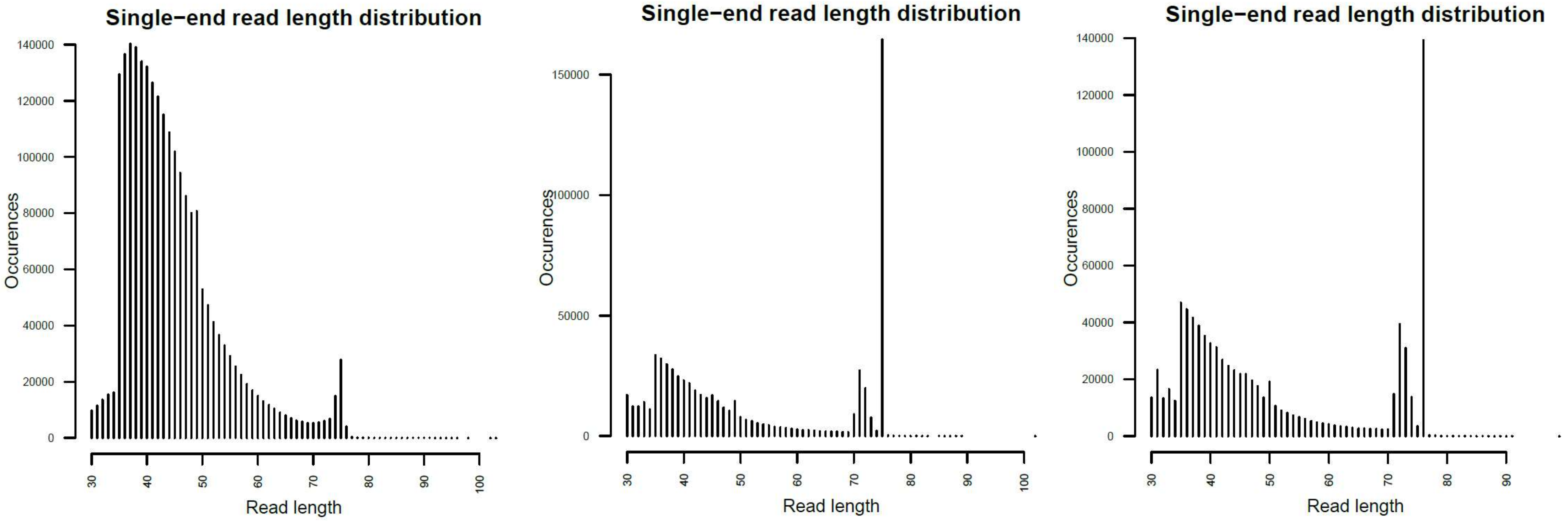


**Fig. S2. Mapped fragment length plots**. From left to right, Par_N16, Par_N9 and Par_N1.

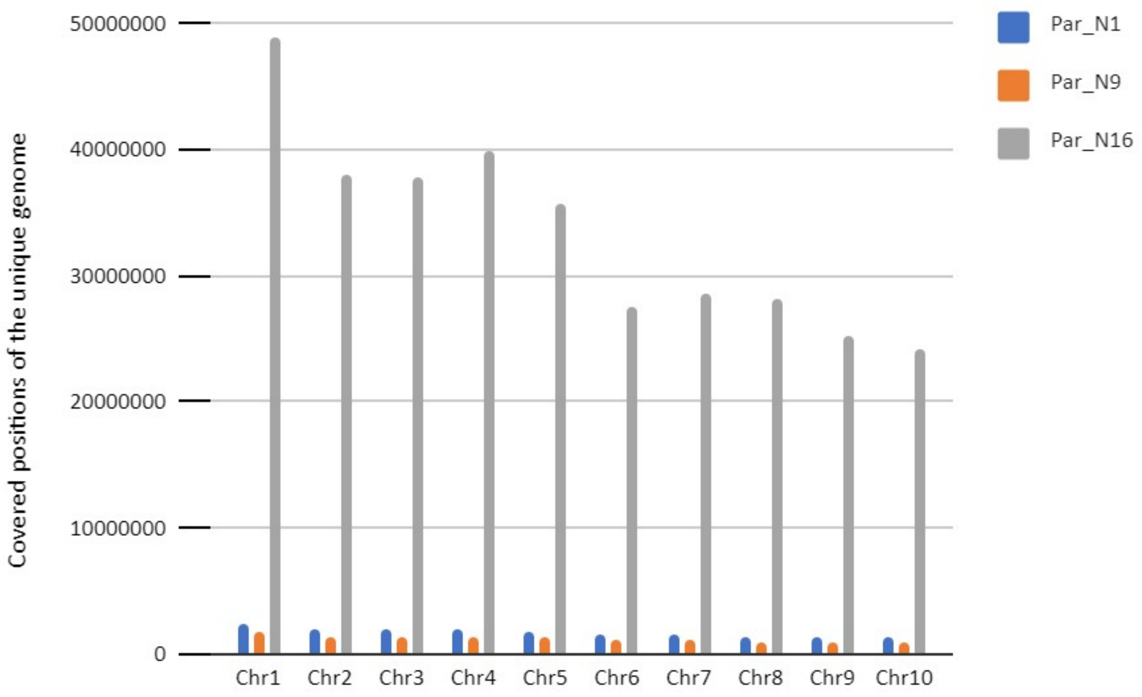


**Fig. S3. Total coverage of the unique genome for the three ancient samples.**

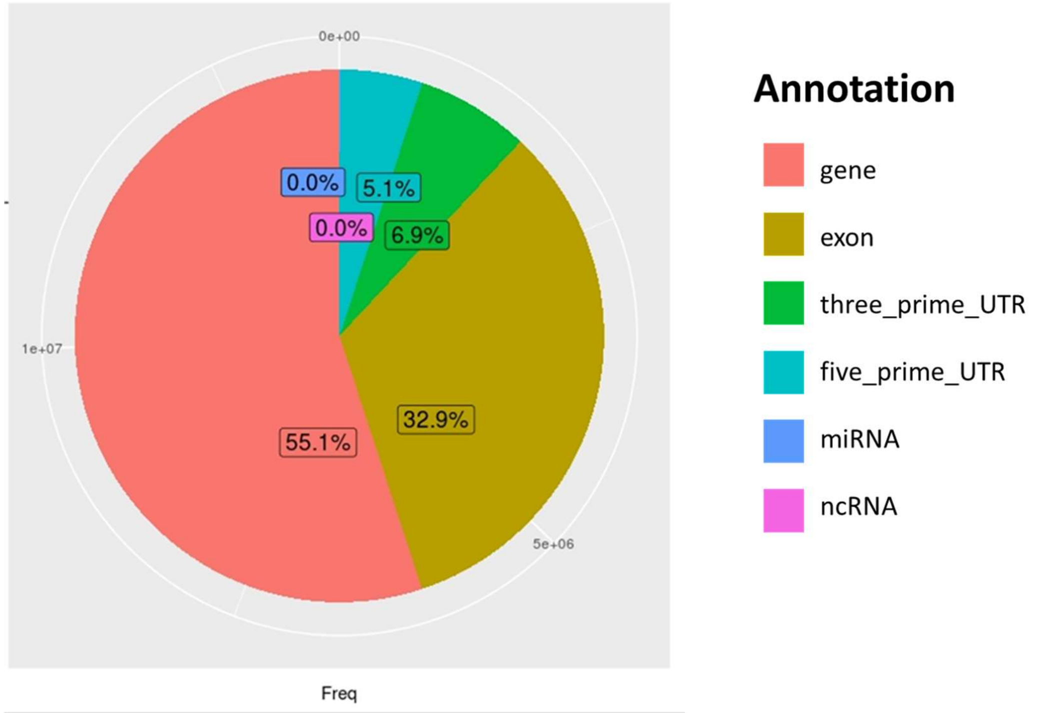


**Fig. S4**. **Distribution of all genomic regions covered by reads from Par_N16**. Percentages of covered sites in Par_N16 with Gramene annotation for genes in orange, exons in brown, 3'-UTR in green, and 5'-UTR in light blue; and miRBase annotation for miRNA in blue, and noncoding RNA in Magenta.

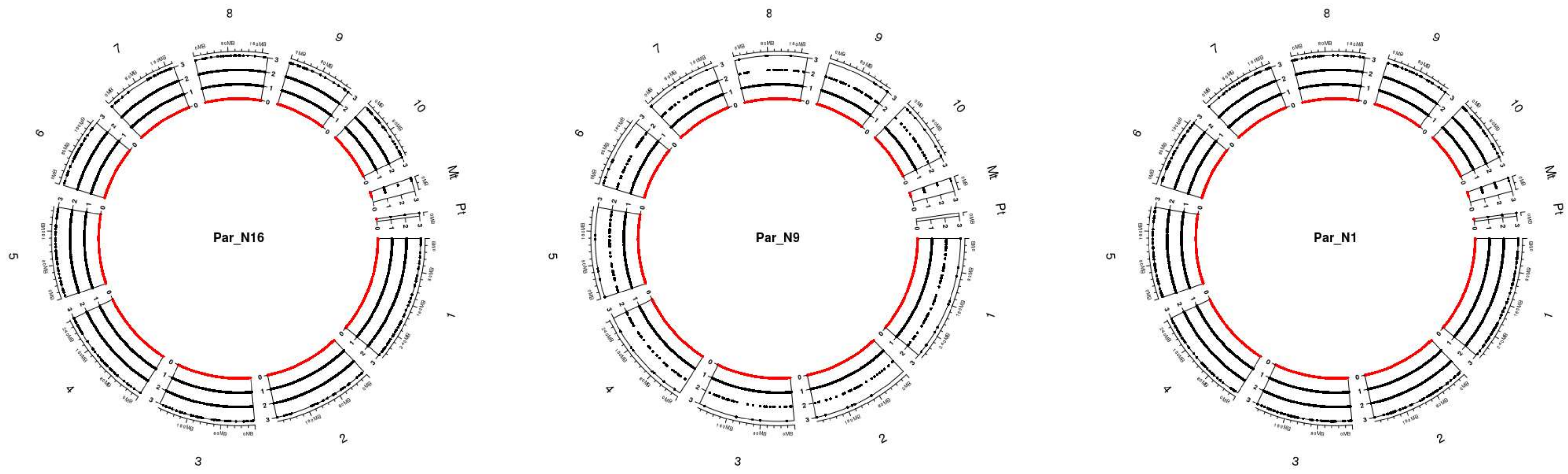


**Fig. S5. Genomic distribution of SNPs for the ancient samples**. The chart includes all ten chromosomes, as well as the chloroplast and mitochondria. Black dots dispersed across chromosomes represent the depth of each SNP scaled on the left of each chromosome, red dots represent sites without coverage.

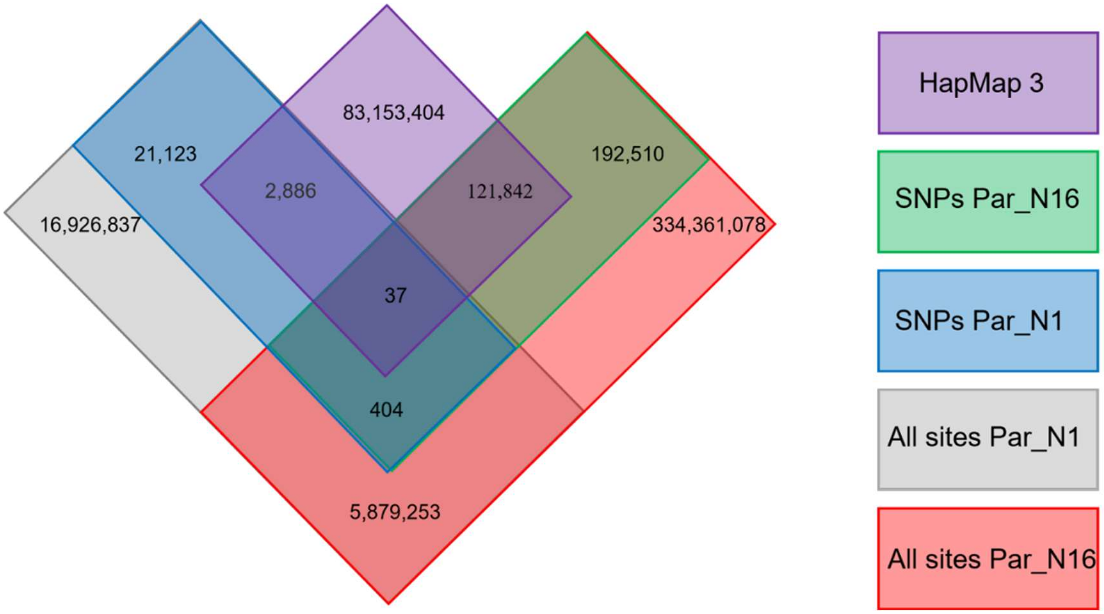


**Fig. S6. Distribution of genotype calls shared between Par_N16, Par_N1, and the HapMap3** (27).

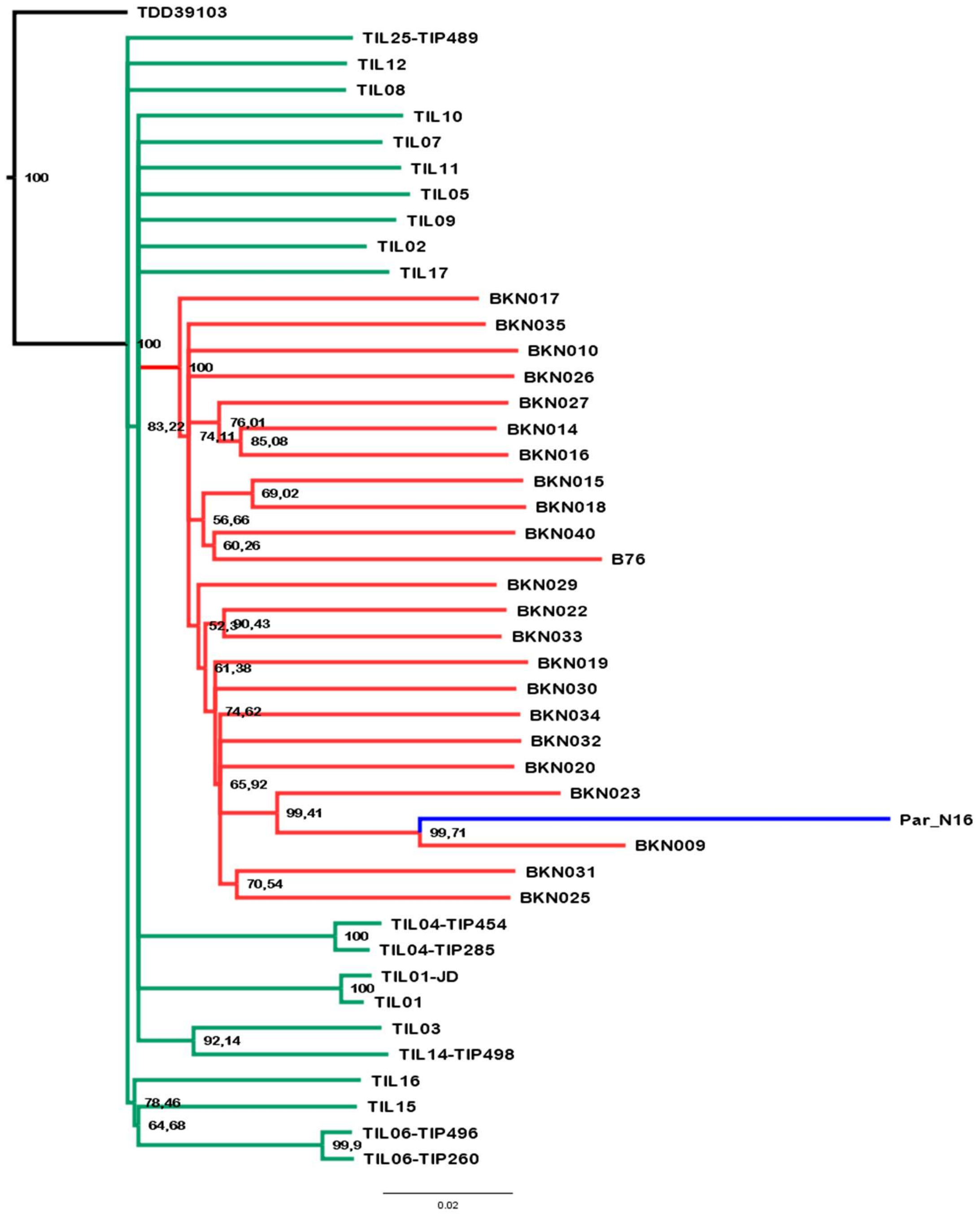


**Fig. S7. Evolutionary relationships between ancient Paredones maize Par_N16 and its wild or cultivated relatives**. Maximum likelihood tree from an alignment of 121,842 genome-wide genotype calls covering non-repetitive regions of the reference maize genome. The teosinte and landrace accessions follow the previously reported nomenclature (27) and are described in Table S5.

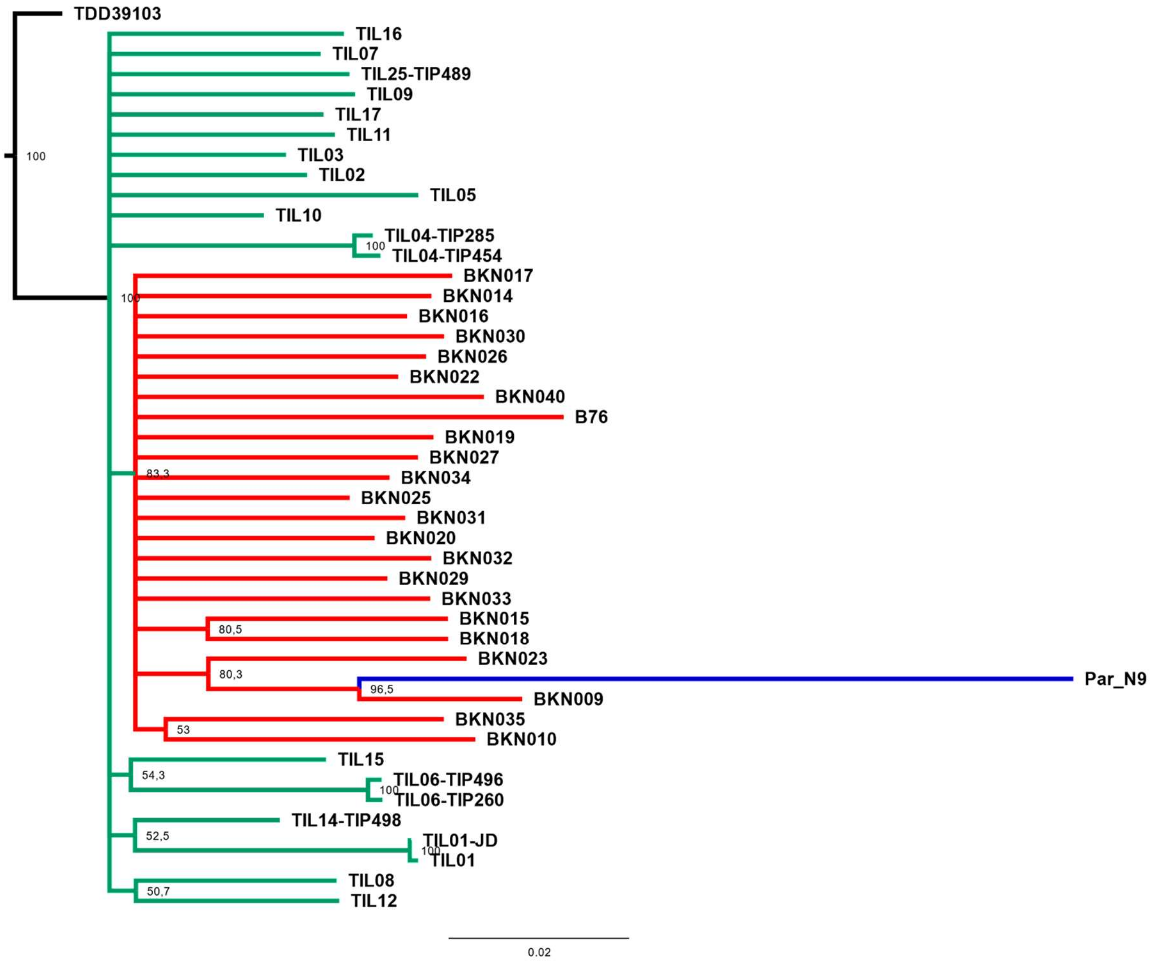


**Fig. S8. Evolutionary relationships between Paredones ancient maize Par_N9 and its wild or cultivated relatives**. Maximum likelihood reconstruction from an alignment of 1,888 genome-wide genotype calls covering non-repetitive regions of the reference maize genome. The teosinte and landrace accessions follow the nomenclature reported in (27) and are described in Table S5.

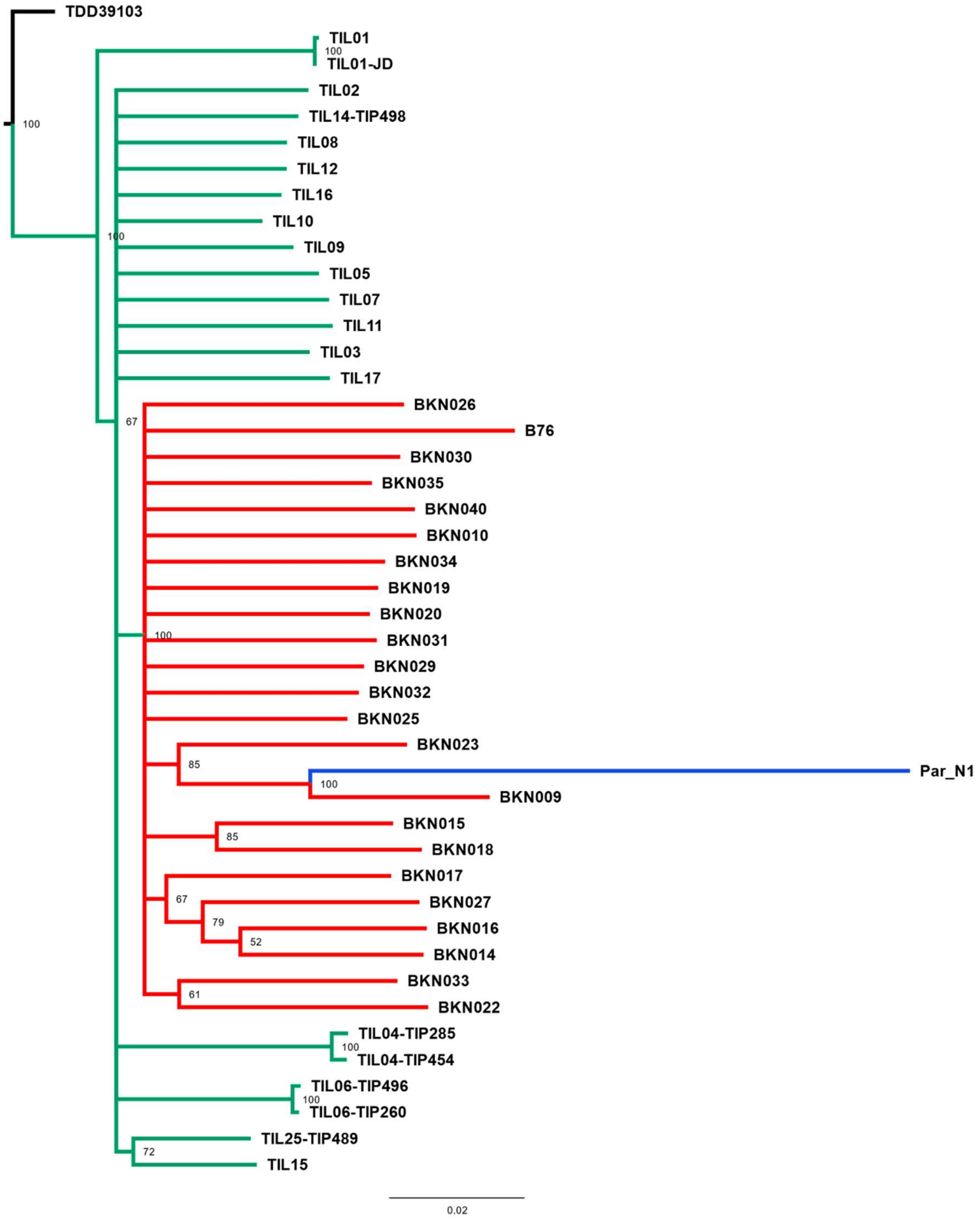


**Fig. S9. Evolutionary relationships between Par_N1 ancient maize and its wild or cultivated relatives**. Maximum likelihood reconstruction from an alignment of 2,886 genome-wide genotype calls covering non-repetitive regions of the reference maize genome. The teosinte and landrace accessions follow the nomenclature reported in (27) and are described in Table S5.

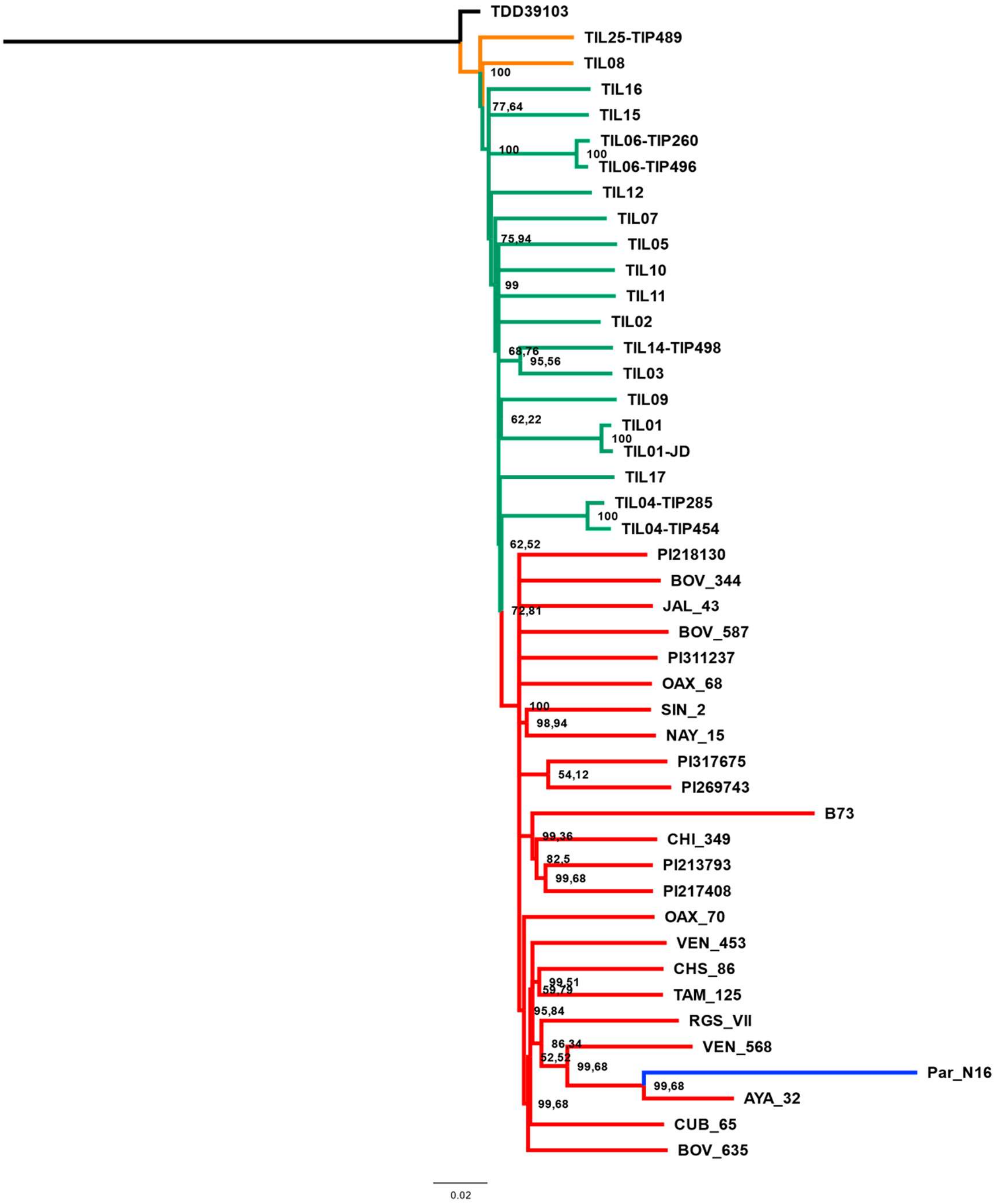


**Fig. S10. Evolutionary relationships between ancient Paredones maize Par_N16 and its wild or cultivated relatives in which only transversions were used**. Maximum likelihood tree from an alignment of 64,118 genome-wide genotype calls covering non-repetitive regions of the reference maize genome. The teosinte and landrace accessions follow the previously reported nomenclature (27) and are described in Table S5.

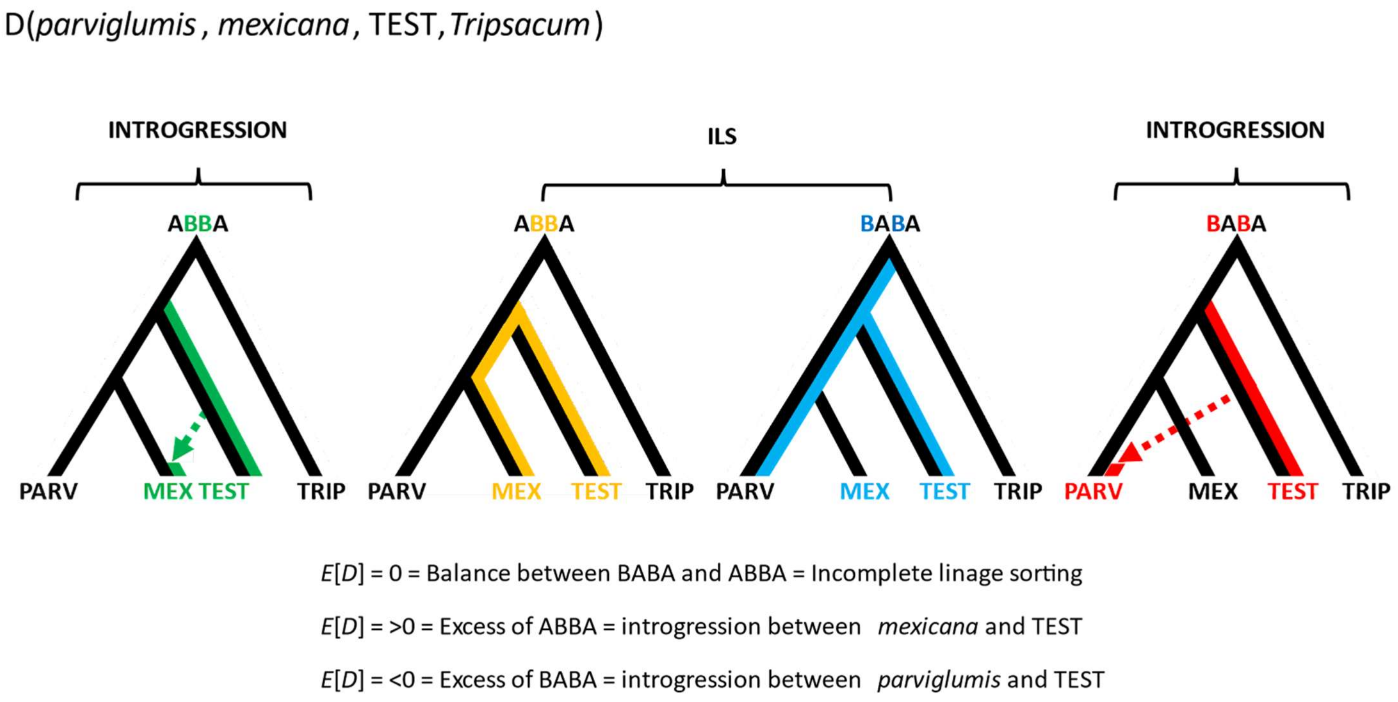


**Fig. S11. Conceptual representation of the hypothesis testing in the form D(*parviglumis*, *mexicana*, TEST, *Tripsacum*)**. The approach makes use of three focal populations, *parviglumis* (PARV), *mexicana* (MEX), and TEST rooted over the outgroup *Tripsacum* (TRIP). Balance between ABBA and BABA sites reflects ILS and/or homogeneous gene flow between TEST and either PARV or MEX. Two other alternatives are: 1) an excess of ABBA resulting in D>0 suggesting introgression between TEST and MEX; 2) an excess of BABA resulting in D<0 suggesting introgression between TEST and PARV.

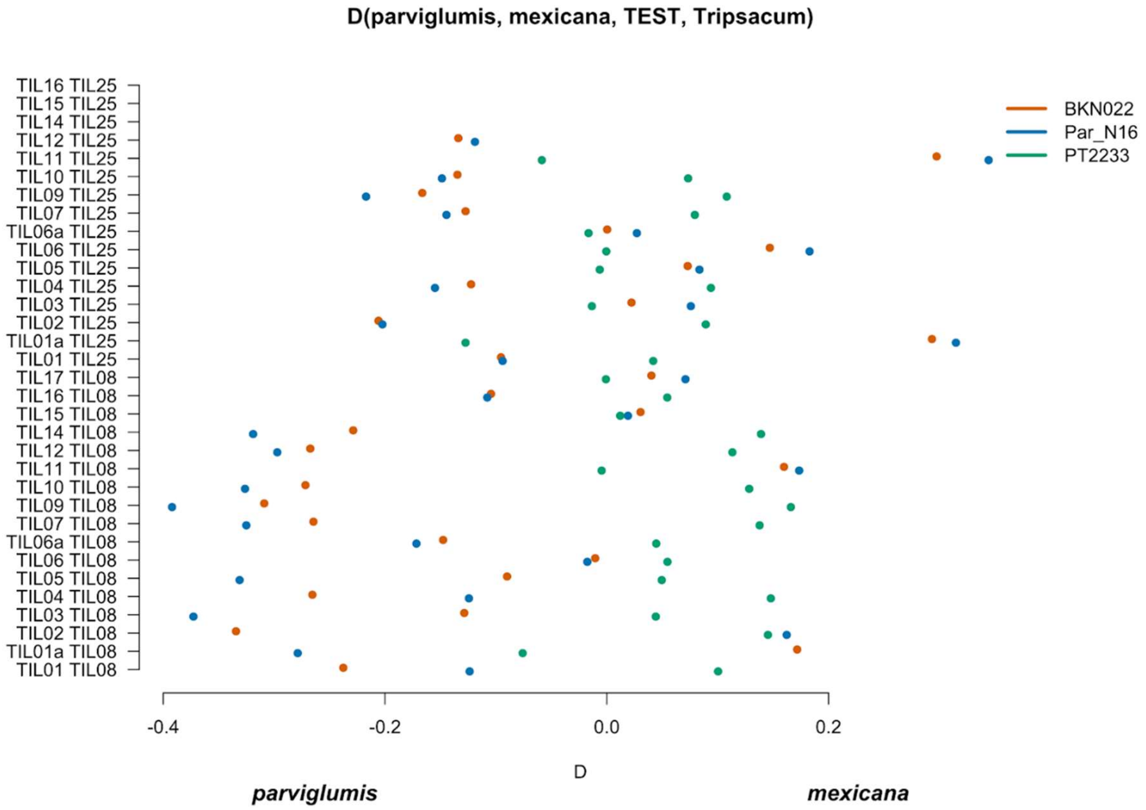


**Fig. S12. Scatterplot of pairwise *mexicana* and *parviglumis* computations of Par_N16, BKN022, and PT2233 D-statistics in the form D(*parviglumis*, *mexicana*, TEST, outgroup)**. Genetic comparison of Par_N16 to teosinte *parviglumis* and *mexicana* accessions. D-statistics were calculated in the form D (*parviglumis*, *mexicana*, TEST, outgroup) (Fig. S11) by comparing 64,118 variant sites shared between Par_N16, *Palomero toluqueño* (PT2233), or *Reventador* (BKN022), and the corresponding SNP variants from teosinte *parviglumis* (TIL01-TIL07, TIL09-17) and two teosinte *mexicana* (TIL25, TIL08) accessions. The graph shows the total number of pairwise comparisons (n=34) (Table S6) yielding a negative (*parviglumis*) or positive (*mexicana*) introgression. Lines in each dot reflect standard deviation calculated from 100 jackknife replicates.

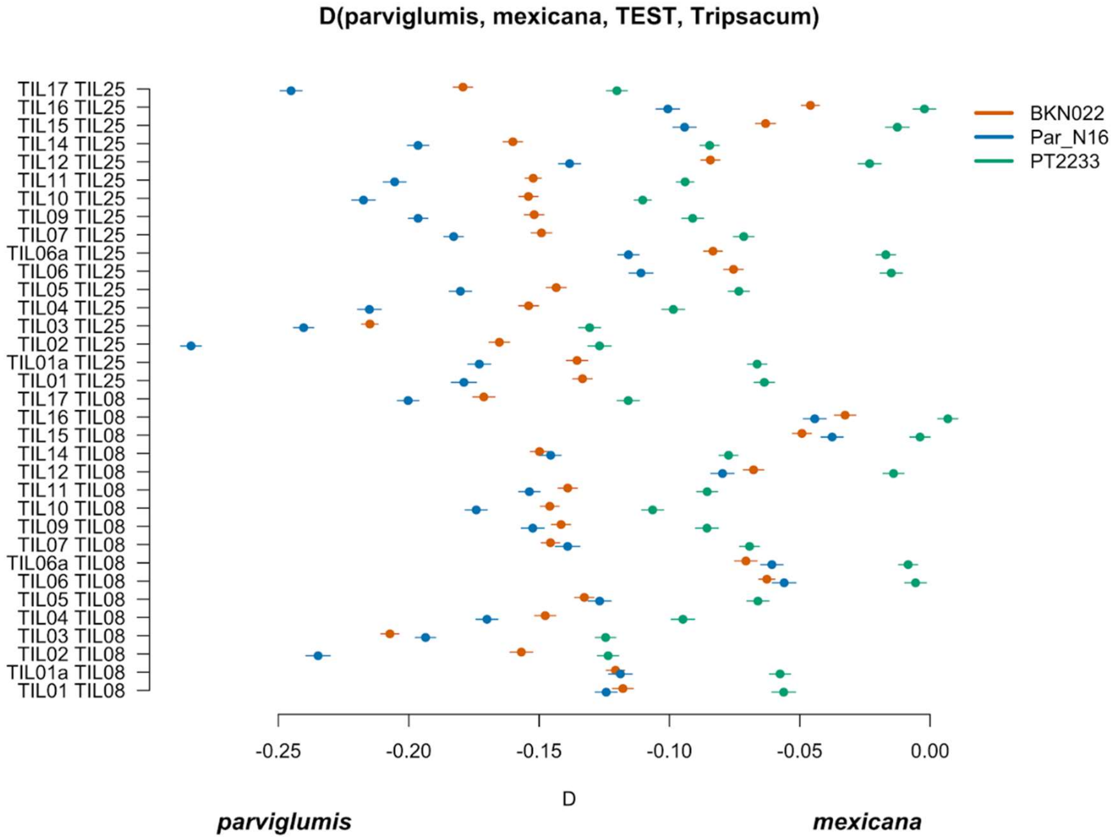


**Fig. S13. Scatterplot of pairwise *mexicana* and *parviglumis* computations of Par_N16, BKN022, and PT2233 D-statistics in the form D(*parviglumis*, *mexicana*, TEST, outgroup) in which only transversions are included**. Genetic comparison of Par_N16 to teosinte *parviglumis* and *mexicana* accessions. D-statistics were calculated in the form D(*parviglumis*, *mexicana*, TEST, outgroup) (Fig. S11) by comparing 64,118 variant sites shared between Par_N16, *Palomero toluqueño* (PT2233), or *Reventador* (NAY15), and the corresponding SNP variants from teosinte *parviglumis* (TIL01-TIL07, TIL09-17) and two teosinte *mexicana* (TIL25, TIL08) accessions. The graph shows the total number of pairwise comparisons (n=34) (Table S7) yielding a negative (*parviglumis*) or positive (*mexicana*) introgression. Lines in each dot reflect standard deviation calculated from 100 jackknife replicates.

D(TEST, *parviglumis* , *mexicana*, *Tripsacum* )

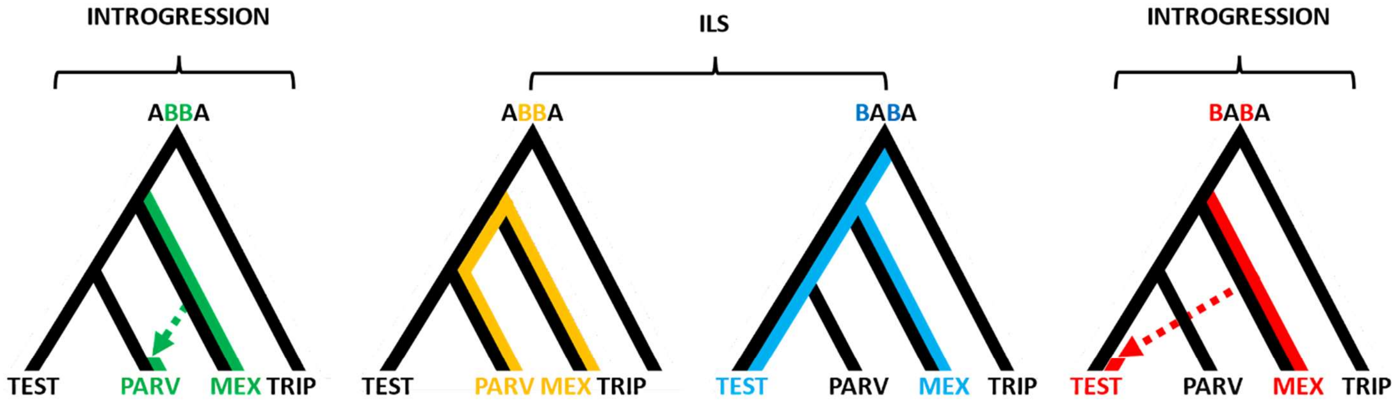


$E[D]$ = 0 = Balance between BABA and ABBA = Incomplete linage sorting

$E[D]$ = >0 = Excess of ABBA = introgression between *mexicana* and *parviglumis*

$E[D]$ = <0 = Excess of BABA = introgression between *mexicana* and TEST

**Fig. S14. Conceptual representation of the hypothesis testing in the form D(TEST, *parviglumis*, *mexicana*, *Tripsacum*)**. The approach makes use of three focal populations; the one being tested (TEST), *parviglumis* (PARV) and *mexicana* (MEX), rooted over the outgroup *Tripsacum* (TRIP). An excess of ABBA sites results in D>0 which suggests that gene flow between *mexicana* and *parviglumis* is higher than the gene flow between *mexicana* and TEST. Meanwhile an excess of BABA sites results in D<0 which suggests that gene flow between *mexicana* and *parviglumis* is lower than the gene flow between *mexicana* and TEST. D=0, given by the same ratio of ABBA and BABA sites reflects ILS and/or the balanced gene flow between the pairs 1) *mexicana* and *parviglumis* and 2) *mexicana* and TEST.

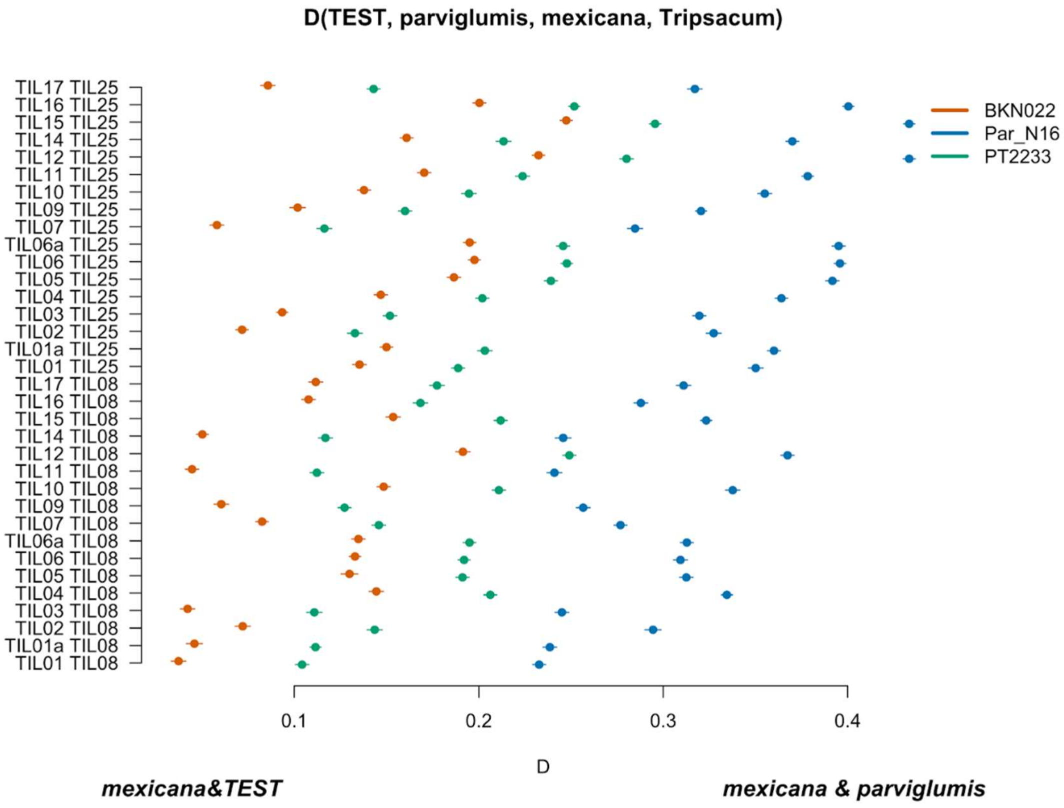


**Fig. S15. Scatterplot of pairwise *mexicana* and *parviglumis* computations of Par_N16, BKN022, and PT2233 D-statistics in the form D(TEST, *parviglumis*, *mexicana*, outgroup) in which only transversions are included**. Genetic comparison of Par_N16 to teosinte *parviglumis* and *mexicana* accessions. D-statistics were calculated in the form D(TEST, *parviglumis*, *mexicana*, outgroup) (Fig. S14) by comparing 64,118 variant sites shared between Par_N16, *Palomero toluqueño* (PT2233), or *Reventador* (BKN022), and the corresponding SNP variants from teosinte *parviglumis* (TIL01-TIL07, TIL09-17) and two teosinte *mexicana* (TIL25, TIL08) accessions. The graph shows the total number of pairwise comparisons (n=34) (Table S9) yielding a negative (*mexicana* & test) or positive (*mexicana* & *parviglumis*) introgression. Lines in each dot reflect standard deviation calculated from 100 jackknife replicates.

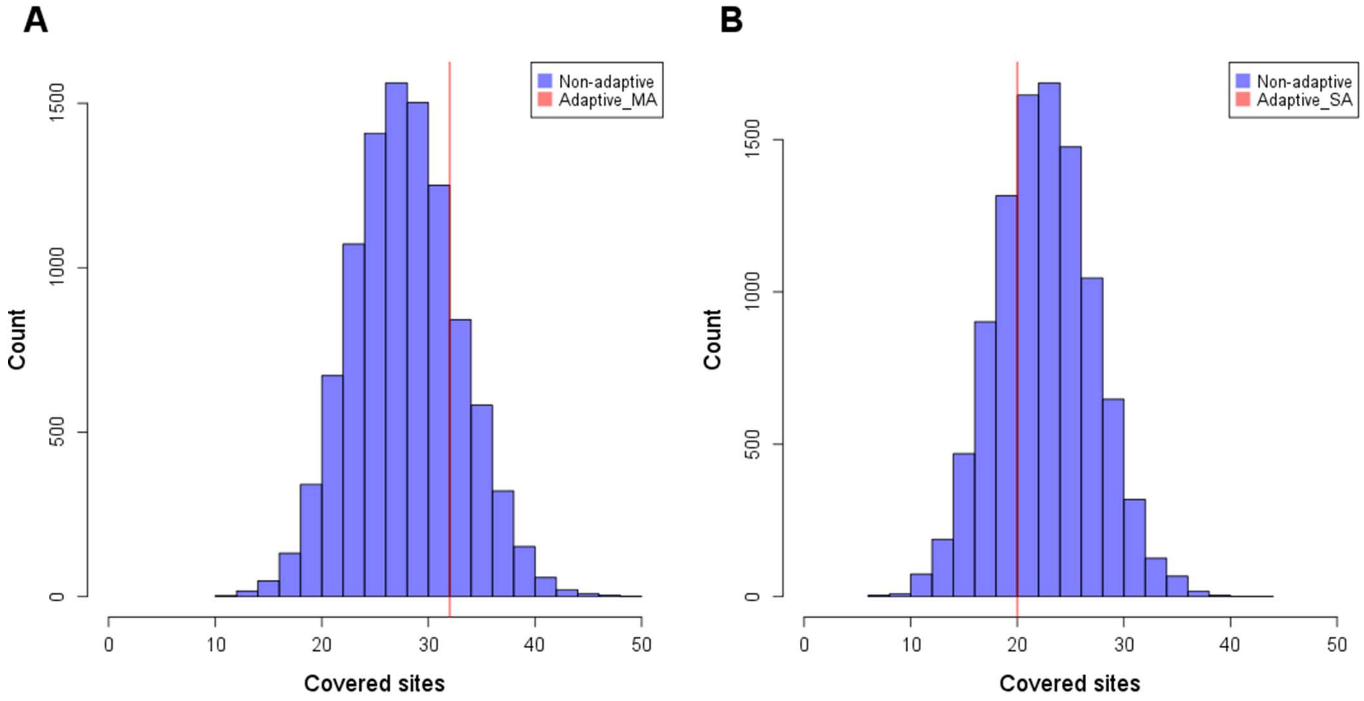


**Fig. S16. Distribution of intersected SNPs between Par_N16 and landraces from Mesoamerica (MA) and South America (SA).** Intersections involved genome-wide non-adaptive SNPs (blue distributions) and SNPs with significant $F_{ST}$ implicated as adaptive (red lines) between Par_N16 and the reference dataset (19) (see Material and Methods). **A**, the genome-wide expected average of covered MA non-adaptive SNPs in Par_N16 is 28.32; the corresponding coverage of adaptive SNPs is 32. **B**, the genome-wide expected average of covered SA non-adaptive SNPs in Par_N16 is 23.02; the corresponding coverage of adaptive SNPs is 20.

**Table S1.** Radiocarbon and calibrated dates of maize specimens from Paredones.

| Sample | Site | Unit \| strata | 14C years BP | 2σ-calibrated age range (BP) | Associated Dating number |
|---|---|---|---|---|---|
| Par-N1(cob) | Paredones | 22\|18 | 5,900±40 | 6,775-6,504 | OS86020 |
| Par-N9 (cob) | Paredones | 20\|4 | 5,300-5,400 | 5,800-5,400 | not assigned* |
| Par-N16 (cob) | Paredones | 20\|5 | 4,849±31 | 5,583–5,324 | AA86936* |
| *Dated stratigraphically | | | | | |
| All data here were published before (see supplementary text) | | | | | |

**Table S2.** Paleogenomic characterization of three ancient maize samples from Paredones.

| | Samples | | |
|---|---|---|---|
| | Par_N1 | Par_N9 | Par_N16 |
| Total number of raw reads | 623,686,255 | 423,856,284 | 851,330,235 |
| Total number of quality sequences | 622,438,882 | 423,472,877 | 850,326,750 |
| Number of sequences mapping to genome | 1,320,284 | 1,034,544 | 15,023,803 |
| Number of sequences mapping to repetitive regions | 441,442 | 351,883 | 5,228,275 |
| Number of sequences mapping to the unique genome | 878,842 | 682,661 | 9,795,586 |
| Total length (Mb) | 52.2 | 40.80 | 471.65 |
| Average read length (bp) | 59.41 | 59.90 | 48.15 |
| Total coverage (Mb) | 16.90 | 12.100 | 334.36 |
| Average quality (Phred) | 31.9 | 32.1 | 34 |
| error rate (mismatches / bases mapped (cigar) | 1,19E-02 | 9,59E-03 | 0,01059832 |

**Table S3.** Total number of unique genomic sites covered at variable depths in ancient Paredones samples.

| Depth | Par_N1 | Par_N9 | Par_N16 |
|---|---|---|---|
| **1** | 15,679,844 | 11,436,116 | 278,622,390 |
| **2** | 1,068,931 | 583,349 | 44,373,018 |
| **3** | 130,670 | 60,892 | 7,898,502 |
| **4** | 24,923 | 11,685 | 1,875,328 |
| **5** | 8,010 | 5,291 | 644,804 |
| **6** | 3,759 | 2,309 | 299,729 |
| **7** | 2,892 | 1,189 | 167,142 |
| **8** | 1,423 | 967 | 103,162 |
| **9** | 1,026 | 837 | 68,272 |
| **10** | 869 | 774 | 47,003 |
| **> 10** | 10,419 | 9341 | 155,805 |

**Table S4.** Number of SNPs and genotype calls recovered from ancient Paredones samples.

| | Par-N1 | Par-N9 | Par-N16 | | Par-N16-transversions |
|---|---|---|---|---|---|
| Total number of SNPs | 21123 | 15554 | 275990 | Total number of SNPs | 275990 |
| Transitions C->T | 7505 | 5766 | 41811 | Transitions C<->T | 76990 |
| Transitions G->A | 7527 | 5617 | 41669 | Transitions G<->A | 76909 |
| INDELS | 609 | 423 | 19790 | INDELS | 19790 |
| Quality SNPs | 5482 | 3748 | 192510 | Quality SNPs | 102302 |
| Genotype calls included in HapMap3 | 2886 | 1888 | 121842 | Genotype calls included in HapMap3 | 64118 |

**Table S5.** Description of all accessions included in the phylogenetic analysis.

| Sample | Line | ID | Race | Species | Class | Country | Accession (source) of origin | Selfing generation |
|---|---|---|---|---|---|---|---|---|
| 1 | TIL01 | TIP-281 | Balsas | *Z. mays ssp. parviglumis* | TIL | Mexico | JSG Y LOS-130 (INIFAP) | S5 |
| 2 | TIL02 | TIP-301 | Balsas | *Z. mays ssp. parviglumis* | TIL | Mexico | JSG Y LOS-119 (INIFAP) | S2 |
| 3 | TIL03 | TIP-282 | Jalisco | *Z. mays ssp. parviglumis* | TIL | Mexico | JSG Y MAS-401 (INIFAP) | S5 |
| 4 | TIL04 | TIP-285 | Balsas | *Z. mays ssp. parviglumis* | TIL | Mexico | 8783 (CIMMYT) | S5 |
| 5 | TIL04 | TIP-454 | Balsas | *Z. mays ssp. parviglumis* | TIL | Mexico | 8783 (CIMMYT) | S5 |
| 6 | TIL05 | TIP-287 | Balsas | *Z. mays ssp. parviglumis* | TIL | Mexico | JSG-197 (INIFAP) | S5 |
| 7 | TIL06 | TIP-260 | Balsas | *Z. mays ssp. parviglumis* | TIL | Mexico | JSG Y LOS-109 (INIFAP) | S3 |
| 8 | TIL06 | TIP-496 | Balsas | *Z. mays ssp. parviglumis* | TIL | Mexico | JSG Y LOS-109 (INIFAP) | S6 |
| 9 | TIL07 | TIP-305 | Balsas | *Z. mays ssp. parviglumis* | TIL | Mexico | JSG-378 (INIFAP) | S4 |
| 10 | TIL08 | TIP-293 | Balsas | *Z. mays ssp. mexicana* | TIL | Mexico | JSG-374 (INIFAP) | S5 |
| 11 | TIL09 | TIP-265 | Balsas | *Z. mays ssp. parviglumis* | TIL | Mexico | JSG Y LOS-161 (INIFAP) | S4 |
| 12 | TIL10 | TIP-304 | Balsas | *Z. mays ssp. parviglumis* | TIL | Mexico | 11355 (CIMMYT) | S4 |
| 13 | TIL11 | TIP-296 | Jalisco | *Z. mays ssp. parviglumis* | TIL | Mexico | JSG Y MAS-264 (INIFAP) | S6 |
| 14 | TIL12 | TIP-267 | Balsas | *Z. mays ssp. parviglumis* | TIL | Mexico | PI566686 (NCRPIS) | S4 |
| 15 | TIL14 | TIP-498 | Jalisco | *Z. mays ssp. parviglumis* | TIL | Mexico | 967 (BFB) | S6 |
| 16 | TIL15 | TIP-276 | Balsas | *Z. mays ssp. parviglumis* | TIL | Mexico | K Site 4 (TAK) | S4 |
| 17 | TIL16 | TIP-309 | Balsas | *Z. mays ssp. parviglumis* | TIL | Mexico | BK Site 4 (GWB) | S4 |
| 18 | TIL17 | TIP-272 | Balsas | *Z. mays ssp. parviglumis* | TIL | Mexico | W Site 6 (HGW) | S4 |
| 19 | TIL25 | TIP-489 | Central Plateau | *Z. mays ssp. mexicana* | TIL | Mexico | 11066 (LMP) | S5 |
| 20 | VEN 568 (NRC) | BKN023 | Araguito | *Z. mays ssp. mays* | LRI | Venezuela | VEN 568 (NRC) | S4 |
| 21 | PI213793 (NCRPIS) | BKN014 | Assiniboine | *Z. mays ssp. mays* | LRI | USA | PI213793 (NCRPIS) | S6 |
| 22 | OAX 68 (INIFAP) | BKN029 | Bolita | *Z. mays ssp. mays* | LRI | Mexico | OAX 68 (INIFAP) | S5 |
| 23 | BOV 635 (NRC) | BKN019 | Cateto | *Z. mays ssp. mays* | LRI | Bolivia | BOV 635 (NRC) | S5 |
| 24 | SIN 2 (INIFAP) | BKN033 | Chapalote | *Z. mays ssp. mays* | LRI | Mexico | SIN 2 (INIFAP) | S5 |
| 25 | CHS 86 (INIFAP) | BKN031 | Comiteco | *Z. mays ssp. mays* | LRI | Mexico | CHS 86 (INIFAP) | S5 |
| 26 | VEN 453 (ICA) | BKN034 | Costeno | *Z. mays ssp. mays* | LRI | Venezuela | VEN 453 (ICA) | S5 |
| 27 | RGS VII (CIMMYT) | BKN032 | Cravo Riogranense | *Z. mays ssp. mays* | LRI | Brazil | RGS VII (CIMMYT) | S5 |

*Continuation Table S5*

| Sample | Line | ID | Race | Species | Class | Country | Accession (source) of origin | Selfing generation |
|---|---|---|---|---|---|---|---|---|
| 28 | CHI 349 (NCGRP) | BKN027 | Cristalino Norteno | *Z. mays ssp. mays* | LRI | Chile | CHI 349 (NCGRP) | S5 |
| 29 | CUB 65 (CIMMYT) | BKN020 | Cuban Flint | *Z. mays ssp. mays* | LRI | Cuba | CUB 65 (CIMMYT) | S5 |
| 30 | PI317675 (NCRPIS) | BKN015 | Havasupai | *Z. mays ssp. mays* | LRI | SW USA | PI317675 (NCRPIS) | S6 |
| 31 | PI311237 (NCRPIS) | BKN040 | Hickory King | *Z. mays ssp. mays* | LRI | USA | PI311237 (NCRPIS) | S7 |
| 32 | PI217408 (NCRPIS) | BKN016 | Longfellow Flint | *Z. mays ssp. mays* | LRI | USA | PI217408 (NCRPIS) | S5 |
| 33 | BOV 344 (ICA) | BKN026 | Pisankalla | *Z. mays ssp. mays* | LRI | Bolivia | BOV 344 (ICA) | S5 |
| 34 | NAY 15 (INIFAP) | BKN022 | Reventador | *Z. mays ssp. mays* | LRI | Mexico | NAY 15 (INIFAP) | S5 |
| 35 | PI218130 (NCRPIS) | BKN017 | Santa Domingo | *Z. mays ssp. mays* | LRI | SW USA | PI218130 (NCRPIS) | S6 |
| 36 | PI269743 (NCRPIS) | BKN018 | Shoe Peg | *Z. mays ssp. mays* | LRI | USA | PI269743 (NCRPIS) | S6 |
| 37 | JAL 43 (CIMMYT) | BKN035 | Tabloncillo | *Z. mays ssp. mays* | LRI | Mexico | JAL 43 (CIMMYT) | S5 |
| 38 | TAM 125 (INIFAP) | BKN025 | Tuxpeno | *Z. mays ssp. mays* | LRI | Mexico | TAM 125 (INIFAP) | S5 |
| 39 | OAX 70 (CIMMYT) | BKN030 | Zapalote Chico | *Z. mays ssp. mays* | LRI | Mexico | OAX 70 (CIMMYT) | S5 |
| 40 | AYA 32 (PCIM) | BKN009 | Chullpi | *Z. mays ssp. mays* | LRI | Peru | AYA 32 (PCIM) | S4 |
| 41 | BOV 587 (NRC) | BKN010 | Poropo | *Z. mays ssp. mays* | LRI | Bolivia | BOV 587 (NRC) | S4 |
| 42 | B73 | B73 | B73 | *Z. mays ssp. mays* | SS | - | B73 | - |
| 43 | TDD39103 | TDD39103 | TDD39103 | *Tripsacum dactyloides* | TRIP | - | TDD39103 | - |

**Table S6.** D-statistic values for each pairwise combination of (*parviglumis*, *mexicana*, TEST, outgroup).

| *parviglumis* | *mexicana* | Test | Outgroup | ABBA sites | BABA sites | D-raw | Z-score | SD-D statistic |
|---|---|---|---|---|---|---|---|---|
| TIL01 | TIL08 | Par_N16 | TDD39103 | 133247 | 170857 | -0.1236748 | 1451.594 | 8.52E-05 |
| TIL01-JD | TIL08 | Par_N16 | TDD39103 | 101992 | 180839 | -0.2787778 | 3929.479 | 7.09E-05 |
| TIL02 | TIL08 | Par_N16 | TDD39103 | 222487 | 160359 | 0.1622793 | 1835.609 | 8.84E-05 |
| TIL03 | TIL08 | Par_N16 | TDD39103 | 87665 | 191894 | -0.3728336 | 5058.11 | 7.37E-05 |
| TIL04-TIP454 | TIL08 | Par_N16 | TDD39103 | 125678 | 161370 | -0.1243416 | 1512.828 | 8.22E-05 |
| TIL05 | TIL08 | Par_N16 | TDD39103 | 91859 | 182832 | -0.331183 | 4308.861 | 7.69E-05 |
| TIL06-TIP260 | TIL08 | Par_N16 | TDD39103 | 147854 | 153140 | -0.01756181 | 9.547868 | 0.001839344 |
| TIL06-TIP496 | TIL08 | Par_N16 | TDD39103 | 121918 | 172468 | -0.1717133 | 95.41205 | 0.001799703 |
| TIL07 | TIL08 | Par_N16 | TDD39103 | 94166 | 184894 | -0.32512 | 4521.422 | 7.19E-05 |
| TIL09 | TIL08 | Par_N16 | TDD39103 | 81798 | 187318 | -0.3920986 | 6183.776 | 6.34E-05 |
| TIL10 | TIL08 | Par_N16 | TDD39103 | 95724 | 188456 | -0.3263143 | 4660.896 | 7.00E-05 |
| TIL11 | TIL08 | Par_N16 | TDD39103 | 197411 | 139036 | 0.1735043 | 1954.53 | 8.88E-05 |
| TIL12 | TIL08 | Par_N16 | TDD39103 | 99219 | 183125 | -0.2971765 | 3621.779 | 8.21E-05 |
| TIL14-TIP498 | TIL08 | Par_N16 | TDD39103 | 92799 | 179733 | -0.3189864 | 4236.013 | 7.53E-05 |
| TIL15 | TIL08 | Par_N16 | TDD39103 | 157410 | 151509 | 0.01910209 | 187.5753 | 0.0001018369 |
| TIL16 | TIL08 | Par_N16 | TDD39103 | 133102 | 165267 | -0.1078028 | 1231.501 | 8.75E-05 |
| TIL17 | TIL08 | Par_N16 | TDD39103 | 142762 | 123847 | 0.07094659 | 844.6917 | 8.40E-05 |
| TIL01 | TIL25-TIP489 | Par_N16 | TDD39103 | 105137 | 126953 | -0.09399802 | 1341.558 | 7.01E-05 |
| TIL01-JD | TIL25-TIP489 | Par_N16 | TDD39103 | 242363 | 126278 | 0.3148999 | 3826.843 | 8.23E-05 |
| TIL02 | TIL25-TIP489 | Par_N16 | TDD39103 | 92475 | 139390 | -0.2023376 | 2851.669 | 7.10E-05 |
| TIL03 | TIL25-TIP489 | Par_N16 | TDD39103 | 142673 | 122566 | 0.0758071 | 820.1362 | 9.24E-05 |
| TIL04-TIP454 | TIL25-TIP489 | Par_N16 | TDD39103 | 94080 | 128592 | -0.1549903 | 2361.327 | 6.56E-05 |
| TIL05 | TIL25-TIP489 | Par_N16 | TDD39103 | 134239 | 113528 | 0.08359063 | 1124.763 | 7.43E-05 |
| TIL06-TIP260 | TIL25-TIP489 | Par_N16 | TDD39103 | 161981 | 111886 | 0.1829173 | 2068.704 | 8.84E-05 |
| TIL06-TIP496 | TIL25-TIP489 | Par_N16 | TDD39103 | 135676 | 128502 | 0.02715593 | 361.1939 | 7.52E-05 |
| TIL07 | TIL25-TIP489 | Par_N16 | TDD39103 | 99677 | 133382 | -0.14462 | 2045.228 | 7.07E-05 |

*Continuation of Table S6*

| *parviglumis* | *mexicana* | Test | Outgroup | ABBA sites | BABA sites | D-raw | Z-score | SD-D statistic |
|---|---|---|---|---|---|---|---|---|
| TIL09 | TIL25-TIP489 | Par_N16 | TDD39103 | 87751 | 136430 | -0.2171415 | 3413.061 | 6.36E-05 |
| TIL10 | TIL25-TIP489 | Par_N16 | TDD39103 | 99240 | 133910 | -0.1487026 | 2196.492 | 6.77E-05 |
| TIL11 | TIL25-TIP489 | Par_N16 | TDD39103 | 218459 | 106525 | 0.3444293 | 4177.508 | 8.24E-05 |
| TIL12 | TIL25-TIP489 | Par_N16 | TDD39103 | 99852 | 126796 | -0.1188804 | 1696.848 | 7.01E-05 |
| TIL14-TIP498 | TIL25-TIP489 | Par_N16 | TDD39103 | 92608 | 123090 | -0.141318 | 2304.143 | 6.13E-05 |
| TIL15 | TIL25-TIP489 | Par_N16 | TDD39103 | 171324 | 109889 | 0.2184643 | 2386.189 | 9.16E-05 |
| TIL16 | TIL25-TIP489 | Par_N16 | TDD39103 | 153102 | 128942 | 0.08566039 | 948.2011 | 9.03E-05 |
| TIL17 | TIL25-TIP489 | Par_N16 | TDD39103 | 103789 | 92362 | 0.05825614 | 1014.304 | 5.74E-05 |
| TIL01 | TIL08 | PT2233 | TDD39103 | 101835 | 83250 | 0.1004133 | 1814.013 | 5.54E-05 |
| TIL01-JD | TIL08 | PT2233 | TDD39103 | 107659 | 125344 | -0.07590031 | 1381.592 | 5.49E-05 |
| TIL02 | TIL08 | PT2233 | TDD39103 | 103209 | 77019 | 0.1453159 | 2824.614 | 5.14E-05 |
| TIL03 | TIL08 | PT2233 | TDD39103 | 99612 | 91180 | 0.04419473 | 917.5232 | 4.82E-05 |
| TIL04-TIP454 | TIL08 | PT2233 | TDD39103 | 101040 | 74997 | 0.1479405 | 3072.645 | 4.81E-05 |
| TIL05 | TIL08 | PT2233 | TDD39103 | 98502 | 89196 | 0.04957964 | 892.1256 | 5.56E-05 |
| TIL06-TIP260 | TIL08 | PT2233 | TDD39103 | 100540 | 90102 | 0.05475184 | 946.8488 | 5.78E-05 |
| TIL06-TIP496 | TIL08 | PT2233 | TDD39103 | 102478 | 93728 | 0.04459599 | 919.5239 | 4.85E-05 |
| TIL07 | TIL08 | PT2233 | TDD39103 | 102104 | 77367 | 0.1378329 | 2885.823 | 4.78E-05 |
| TIL09 | TIL08 | PT2233 | TDD39103 | 100240 | 71693 | 0.1660356 | 3672.995 | 4.52E-05 |
| TIL10 | TIL08 | PT2233 | TDD39103 | 102469 | 79138 | 0.1284697 | 2694.865 | 4.77E-05 |
| TIL11 | TIL08 | PT2233 | TDD39103 | 100595 | 101537 | -0.004660321 | 77.23002 | 6.03E-05 |
| TIL12 | TIL08 | PT2233 | TDD39103 | 102727 | 81824 | 0.1132641 | 2169.879 | 5.22E-05 |
| TIL14-TIP498 | TIL08 | PT2233 | TDD39103 | 99344 | 75080 | 0.1391093 | 2631.956 | 5.29E-05 |
| TIL15 | TIL08 | PT2233 | TDD39103 | 99655 | 97276 | 0.01208037 | 220.1007 | 5.49E-05 |
| TIL16 | TIL08 | PT2233 | TDD39103 | 100973 | 90526 | 0.05455381 | 1054.854 | 5.17E-05 |
| TIL17 | TIL08 | PT2233 | TDD39103 | 90264 | 90402 | -0.0007638405 | 13.84347 | 5.52E-05 |
| TIL01 | TIL25-TIP489 | PT2233 | TDD39103 | 86952 | 79967 | 0.04184664 | 795.8017 | 5.26E-05 |
| TIL01-JD | TIL25-TIP489 | PT2233 | TDD39103 | 99068 | 128000 | -0.1274156 | 2304.53 | 5.53E-05 |
| TIL02 | TIL25-TIP489 | PT2233 | TDD39103 | 90156 | 75374 | 0.08930103 | 1677.873 | 5.32E-05 |
| TIL03 | TIL25-TIP489 | PT2233 | TDD39103 | 89313 | 91746 | -0.01343761 | 267.6182 | 5.02E-05 |
| TIL04-TIP454 | TIL25-TIP489 | PT2233 | TDD39103 | 87220 | 72240 | 0.09394205 | 1888.075 | 4.98E-05 |

*Continuation of Table S6*

| *parviglumis* | *mexicana* | Test | Outgroup | ABBA sites | BABA sites | D-raw | Z-score | SD-D statistic |
|---|---|---|---|---|---|---|---|---|
| TIL05 | TIL25-TIP489 | PT2233 | TDD39103 | 86256 | 87357 | -0.006341691 | 120.0282 | 5.28E-05 |
| TIL06-TIP260 | TIL25-TIP489 | PT2233 | TDD39103 | 88944 | 89022 | -0.000438286 | 7.841583 | 5.59E-05 |
| TIL06-TIP496 | TIL25-TIP489 | PT2233 | TDD39103 | 91668 | 94731 | -0.01643249 | 351.7436 | 4.67E-05 |
| TIL07 | TIL25-TIP489 | PT2233 | TDD39103 | 89490 | 76325 | 0.07939571 | 1599.028 | 4.97E-05 |
| TIL09 | TIL25-TIP489 | PT2233 | TDD39103 | 88429 | 71168 | 0.1081537 | 2264.205 | 4.78E-05 |
| TIL10 | TIL25-TIP489 | PT2233 | TDD39103 | 87469 | 75512 | 0.07336438 | 1566.824 | 4.68E-05 |
| TIL01 | TIL08 | BKN022 | TDD39103 | 113212 | 183809 | -0.2376835 | 132.6043 | 0.001792427 |
| TIL01-JD | TIL08 | BKN022 | TDD39103 | 230720 | 163092 | 0.1717266 | 106.8575 | 0.001607061 |
| TIL02 | TIL08 | BKN022 | TDD39103 | 100092 | 200673 | -0.3344172 | 193.1228 | 0.00173163 |
| TIL03 | TIL08 | BKN022 | TDD39103 | 132536 | 171633 | -0.1285371 | 72.95316 | 0.001761913 |
| TIL04-TIP454 | TIL08 | BKN022 | TDD39103 | 105656 | 182060 | -0.2655535 | 148.0532 | 0.001793636 |
| TIL05 | TIL08 | BKN022 | TDD39103 | 135235 | 161955 | -0.08990881 | 50.23206 | 0.001789869 |
| TIL06-TIP260 | TIL08 | BKN022 | TDD39103 | 155700 | 158994 | -0.01046731 | 5.612727 | 0.001864924 |
| TIL06-TIP496 | TIL08 | BKN022 | TDD39103 | 131518 | 177075 | -0.1476281 | 81.46008 | 0.001812275 |
| TIL07 | TIL08 | BKN022 | TDD39103 | 108540 | 186640 | -0.2645843 | 150.6103 | 0.001756748 |
| TIL09 | TIL08 | BKN022 | TDD39103 | 97039 | 183820 | -0.3089842 | 172.6957 | 0.001789183 |
| TIL10 | TIL08 | BKN022 | TDD39103 | 108545 | 189579 | -0.2718131 | 158.6396 | 0.0017134 |
| TIL11 | TIL08 | BKN022 | TDD39103 | 201207 | 145750 | 0.1598383 | 97.97877 | 0.001631356 |
| TIL12 | TIL08 | BKN022 | TDD39103 | 110503 | 191185 | -0.2674352 | 150.5364 | 0.001776549 |
| TIL14-TIP498 | TIL08 | BKN022 | TDD39103 | 108232 | 172427 | -0.2287295 | 125.3859 | 0.001824204 |
| TIL15 | TIL08 | BKN022 | TDD39103 | 165052 | 155301 | 0.0304383 | 17.76504 | 0.001713382 |
| TIL16 | TIL08 | BKN022 | TDD39103 | 140143 | 172830 | -0.1044403 | 60.53167 | 0.001725383 |
| TIL17 | TIL08 | BKN022 | TDD39103 | 147975 | 136538 | 0.04019851 | 22.11122 | 0.001818014 |
| TIL01 | TIL25-TIP489 | BKN022 | TDD39103 | 114346 | 138462 | -0.09539255 | 46.82806 | 0.002037081 |
| TIL01-JD | TIL25-TIP489 | BKN022 | TDD39103 | 245287 | 134036 | 0.2932883 | 184.4768 | 0.001589838 |
| TIL02 | TIL25-TIP489 | BKN022 | TDD39103 | 102869 | 156206 | -0.2058747 | 110.8628 | 0.001857024 |
| TIL03 | TIL25-TIP489 | BKN022 | TDD39103 | 143910 | 137632 | 0.02229863 | 11.91879 | 0.00187088 |
| TIL04-TIP454 | TIL25-TIP489 | BKN022 | TDD39103 | 107037 | 136893 | -0.1223958 | 59.8654 | 0.002044516 |
| TIL05 | TIL25-TIP489 | BKN022 | TDD39103 | 142463 | 123092 | 0.07294534 | 39.08395 | 0.001866376 |
| TIL06-TIP260 | TIL25-TIP489 | BKN022 | TDD39103 | 166720 | 123977 | 0.1470363 | 82.74616 | 0.001776956 |

*Continuation of Table S6*

| *parviglumis* | *mexicana* | Test | Outgroup | ABBA sites | BABA sites | D-raw | Z-score | SD-D statistic |
|---|---|---|---|---|---|---|---|---|
| TIL06-TIP496 | TIL25-TIP489 | BKN022 | TDD39103 | 142316 | 142215 | 0.0003549701 | 0.1948321 | 0.001821929 |
| TIL07 | TIL25-TIP489 | BKN022 | TDD39103 | 111431 | 143960 | -0.1273694 | 65.02995 | 0.001958627 |
| TIL09 | TIL25-TIP489 | BKN022 | TDD39103 | 101182 | 141590 | -0.1664442 | 86.413 | 0.001926148 |
| TIL10 | TIL25-TIP489 | BKN022 | TDD39103 | 109529 | 143628 | -0.1346951 | 70.54111 | 0.001909455 |
| TIL11 | TIL25-TIP489 | BKN022 | TDD39103 | 218566 | 118343 | 0.297478 | 186.829 | 0.001592248 |
| TIL12 | TIL25-TIP489 | BKN022 | TDD39103 | 109350 | 143166 | -0.1339163 | 69.20308 | 0.00193512 |
| TIL14-TIP498 | TIL25-TIP489 | BKN022 | TDD39103 | 106853 | 126522 | -0.08428066 | 1337.487 | 6.30E-05 |
| TIL15 | TIL25-TIP489 | BKN022 | TDD39103 | 174654 | 120483 | 0.1835453 | 2140.452 | 8.58E-05 |
| TIL16 | TIL25-TIP489 | BKN022 | TDD39103 | 156348 | 143022 | 0.04451348 | 542.2696 | 8.21E-05 |

**Table S7.** D-statistic values for each pairwise combination of (*parviglumis*, *mexicana*, TEST, outgroup) including only transversions.

| *parviglumis* | *mexicana* | Test | Outgroup | ABBA sites | BABA sites | D-raw | Z-score | SD-D statistic |
|---|---|---|---|---|---|---|---|---|
| TIL01 | TIL08 | Par_N16 | TDD39103 | 23899 | 30682 | -0.124274 | 29.47079 | 0.004216854 |
| TIL01-JD | TIL08 | Par_N16 | TDD39103 | 24663 | 31313 | -0.1188009 | 26.21163 | 0.004532375 |
| TIL02 | TIL08 | Par_N16 | TDD39103 | 18079 | 29173 | -0.2347837 | 50.56658 | 0.00464306 |
| TIL03 | TIL08 | Par_N16 | TDD39103 | 23000 | 34042 | -0.1935767 | 49.46089 | 0.003913732 |
| TIL04-TIP454 | TIL08 | Par_N16 | TDD39103 | 21569 | 30405 | -0.1700081 | 40.14312 | 0.004235049 |
| TIL05 | TIL08 | Par_N16 | TDD39103 | 23650 | 30518 | -0.1267907 | 28.65204 | 0.00442519 |
| TIL06-TIP260 | TIL08 | Par_N16 | TDD39103 | 23933 | 26772 | -0.05599053 | 12.39952 | 0.004515541 |
| TIL06-TIP496 | TIL08 | Par_N16 | TDD39103 | 23427 | 26455 | -0.06070326 | 14.00943 | 0.004333029 |
| TIL07 | TIL08 | Par_N16 | TDD39103 | 22781 | 30140 | -0.1390563 | 29.4367 | 0.00472391 |
| TIL09 | TIL08 | Par_N16 | TDD39103 | 23646 | 32151 | -0.1524275 | 34.70303 | 0.004392341 |
| TIL10 | TIL08 | Par_N16 | TDD39103 | 22283 | 31679 | -0.1741225 | 40.85013 | 0.004262472 |
| TIL11 | TIL08 | Par_N16 | TDD39103 | 23950 | 32652 | -0.1537402 | 37.45568 | 0.004104588 |
| TIL12 | TIL08 | Par_N16 | TDD39103 | 21798 | 25571 | -0.07965125 | 17.8432 | 0.004463956 |
| TIL14-TIP498 | TIL08 | Par_N16 | TDD39103 | 23764 | 31861 | -0.145564 | 36.19816 | 0.004021311 |
| TIL15 | TIL08 | Par_N16 | TDD39103 | 24928 | 26875 | -0.0375847 | 8.833977 | 0.004254561 |
| TIL16 | TIL08 | Par_N16 | TDD39103 | 23929 | 26145 | -0.0442545 | 10.08735 | 0.00438713 |
| TIL17 | TIL08 | Par_N16 | TDD39103 | 21301 | 31969 | -0.2002628 | 48.40805 | 0.004136973 |
| TIL01 | TIL25-TIP489 | Par_N16 | TDD39103 | 22070 | 31684 | -0.1788518 | 37.06851 | 0.004824899 |
| TIL01-JD | TIL25-TIP489 | Par_N16 | TDD39103 | 22788 | 32319 | -0.1729544 | 39.13436 | 0.004419504 |
| TIL02 | TIL25-TIP489 | Par_N16 | TDD39103 | 17717 | 31739 | -0.2835247 | 70.59782 | 0.004016055 |
| TIL03 | TIL25-TIP489 | Par_N16 | TDD39103 | 22054 | 36008 | -0.2403293 | 61.59418 | 0.003901819 |
| TIL04-TIP454 | TIL25-TIP489 | Par_N16 | TDD39103 | 21390 | 33114 | -0.2151035 | 47.23022 | 0.004554361 |
| TIL05 | TIL25-TIP489 | Par_N16 | TDD39103 | 22242 | 32019 | -0.1801847 | 41.9849 | 0.004291654 |
| TIL06-TIP260 | TIL25-TIP489 | Par_N16 | TDD39103 | 22530 | 28152 | -0.110927 | 24.16857 | 0.00458972 |
| TIL06-TIP496 | TIL25-TIP489 | Par_N16 | TDD39103 | 22126 | 27917 | -0.1157205 | 28.2432 | 0.004097287 |
| TIL07 | TIL25-TIP489 | Par_N16 | TDD39103 | 22702 | 32857 | -0.1827787 | 48.95903 | 0.003733298 |

*Continuation of Table S7*

| *parviglumis* | *mexicana* | Test | Outgroup | ABBA sites | BABA sites | D-raw | Z-score | SD-D statistic |
|---|---|---|---|---|---|---|---|---|
| TIL09 | TIL25-TIP489 | Par_N16 | TDD39103 | 22931 | 34142 | -0.1964326 | 52.58948 | 0.003735208 |
| TIL10 | TIL25-TIP489 | Par_N16 | TDD39103 | 22179 | 34499 | -0.2173683 | 48.14574 | 0.004514798 |
| TIL11 | TIL25-TIP489 | Par_N16 | TDD39103 | 22477 | 34099 | -0.2054228 | 46.77688 | 0.004391546 |
| TIL12 | TIL25-TIP489 | Par_N16 | TDD39103 | 20299 | 26815 | -0.1383028 | 32.61333 | 0.004240685 |
| TIL14-TIP498 | TIL25-TIP489 | Par_N16 | TDD39103 | 22477 | 33467 | -0.1964464 | 47.81359 | 0.00410859 |
| TIL15 | TIL25-TIP489 | Par_N16 | TDD39103 | 22965 | 27741 | -0.09419004 | 21.15305 | 0.004452788 |
| TIL16 | TIL25-TIP489 | Par_N16 | TDD39103 | 22049 | 26983 | -0.1006282 | 22.1901 | 0.004534822 |
| TIL17 | TIL25-TIP489 | Par_N16 | TDD39103 | 21044 | 34713 | -0.2451531 | 57.99813 | 0.004226913 |
| TIL01 | TIL08 | PT2233 | TDD39103 | 30154 | 33743 | -0.05616852 | 12.39773 | 0.004530548 |
| TIL01-JD | TIL08 | PT2233 | TDD39103 | 31004 | 34791 | -0.05755757 | 14.26732 | 0.004034224 |
| TIL02 | TIL08 | PT2233 | TDD39103 | 24276 | 31119 | -0.123531 | 30.19098 | 0.004091653 |
| TIL03 | TIL08 | PT2233 | TDD39103 | 29429 | 37798 | -0.1244887 | 31.66375 | 0.003931584 |
| TIL04-TIP454 | TIL08 | PT2233 | TDD39103 | 27659 | 33454 | -0.09482434 | 20.91208 | 0.004534428 |
| TIL05 | TIL08 | PT2233 | TDD39103 | 29717 | 33921 | -0.06606116 | 15.53563 | 0.004252235 |
| TIL06-TIP260 | TIL08 | PT2233 | TDD39103 | 29740 | 30075 | -0.005600602 | 1.353725 | 0.004137178 |
| TIL06-TIP496 | TIL08 | PT2233 | TDD39103 | 29135 | 29631 | -0.008440255 | 2.298614 | 0.003671888 |
| TIL07 | TIL08 | PT2233 | TDD39103 | 29045 | 33369 | -0.06927933 | 18.15405 | 0.003816191 |
| TIL09 | TIL08 | PT2233 | TDD39103 | 29943 | 35550 | -0.0856122 | 19.54822 | 0.004379539 |
| TIL10 | TIL08 | PT2233 | TDD39103 | 28477 | 35262 | -0.1064497 | 25.3051 | 0.004206651 |
| TIL11 | TIL08 | PT2233 | TDD39103 | 30355 | 36033 | -0.0855275 | 21.24596 | 0.004025589 |
| TIL12 | TIL08 | PT2233 | TDD39103 | 27349 | 28128 | -0.01404186 | 3.485315 | 0.004028862 |
| TIL14-TIP498 | TIL08 | PT2233 | TDD39103 | 30189 | 35250 | -0.0773392 | 21.18885 | 0.003649996 |
| TIL15 | TIL08 | PT2233 | TDD39103 | 30413 | 30650 | -0.003881237 | 0.9986657 | 0.003886423 |
| TIL16 | TIL08 | PT2233 | TDD39103 | 29699 | 29299 | 0.006779891 | 1.738047 | 0.003900866 |
| TIL17 | TIL08 | PT2233 | TDD39103 | 27538 | 34752 | -0.1158131 | 26.97966 | 0.004292609 |
| TIL01 | TIL25-TIP489 | PT2233 | TDD39103 | 30159 | 34256 | -0.0636032 | 16.28916 | 0.003904632 |
| TIL01-JD | TIL25-TIP489 | PT2233 | TDD39103 | 30866 | 35254 | -0.06636419 | 17.69144 | 0.003751204 |
| TIL02 | TIL25-TIP489 | PT2233 | TDD39103 | 25818 | 33318 | -0.1268263 | 28.1771 | 0.004501042 |
| TIL03 | TIL25-TIP489 | PT2233 | TDD39103 | 30258 | 39349 | -0.1306047 | 30.84884 | 0.004233699 |
| TIL04-TIP454 | TIL25-TIP489 | PT2233 | TDD39103 | 29340 | 35753 | -0.09852058 | 22.41358 | 0.004395575 |

*Continuation of Table S7*

| *parviglumis* | *mexicana* | Test | Outgroup | ABBA sites | BABA sites | D-raw | Z-score | SD-D statistic |
|---|---|---|---|---|---|---|---|---|
| TIL05 | TIL25-TIP489 | PT2233 | TDD39103 | 30097 | 34865 | -0.07339676 | 18.11266 | 0.004052235 |
| TIL06-TIP260 | TIL25-TIP489 | PT2233 | TDD39103 | 30285 | 31201 | -0.0148977 | 3.475785 | 0.00428614 |
| TIL06-TIP496 | TIL25-TIP489 | PT2233 | TDD39103 | 29806 | 30837 | -0.01700114 | 4.531805 | 0.003751516 |
| TIL07 | TIL25-TIP489 | PT2233 | TDD39103 | 30910 | 35670 | -0.07149294 | 18.02631 | 0.003966032 |
| TIL09 | TIL25-TIP489 | PT2233 | TDD39103 | 30948 | 37151 | -0.09108797 | 21.73673 | 0.00419051 |
| TIL10 | TIL25-TIP489 | PT2233 | TDD39103 | 30110 | 37572 | -0.1102509 | 33.64911 | 0.003276487 |
| TIL11 | TIL25-TIP489 | PT2233 | TDD39103 | 30527 | 36861 | -0.093993 | 27.67844 | 0.003395893 |
| TIL12 | TIL25-TIP489 | PT2233 | TDD39103 | 27785 | 29105 | -0.02320267 | 5.228515 | 0.004437717 |
| TIL14-TIP498 | TIL25-TIP489 | PT2233 | TDD39103 | 30681 | 36351 | -0.08458647 | 22.95687 | 0.003684582 |
| TIL15 | TIL25-TIP489 | PT2233 | TDD39103 | 30264 | 31035 | -0.01257769 | 2.763191 | 0.004551872 |
| TIL16 | TIL25-TIP489 | PT2233 | TDD39103 | 29586 | 29715 | -0.002175343 | 0.4977587 | 0.004370276 |
| TIL17 | TIL25-TIP489 | PT2233 | TDD39103 | 29294 | 37296 | -0.1201682 | 29.9125 | 0.004017323 |
| TIL01 | TIL08 | BKN022 | TDD39103 | 31935 | 40468 | -0.1178542 | 29.35414 | 0.00401491 |
| TIL01-JD | TIL08 | BKN022 | TDD39103 | 32703 | 41681 | -0.120698 | 33.91904 | 0.003558415 |
| TIL02 | TIL08 | BKN022 | TDD39103 | 26125 | 35842 | -0.1568093 | 36.08191 | 0.004345925 |
| TIL03 | TIL08 | BKN022 | TDD39103 | 30644 | 46664 | -0.2072231 | 59.86631 | 0.00346143 |
| TIL04-TIP454 | TIL08 | BKN022 | TDD39103 | 29362 | 39535 | -0.1476552 | 36.48487 | 0.004047025 |
| TIL05 | TIL08 | BKN022 | TDD39103 | 31263 | 40826 | -0.1326555 | 35.65501 | 0.003720528 |
| TIL06-TIP260 | TIL08 | BKN022 | TDD39103 | 31507 | 35718 | -0.06264039 | 20.17036 | 0.003105566 |
| TIL06-TIP496 | TIL08 | BKN022 | TDD39103 | 30840 | 35531 | -0.07067846 | 16.46917 | 0.004291562 |
| TIL07 | TIL08 | BKN022 | TDD39103 | 30506 | 40907 | -0.1456458 | 40.89704 | 0.003561279 |
| TIL09 | TIL08 | BKN022 | TDD39103 | 32023 | 42586 | -0.1415781 | 38.38372 | 0.003688494 |
| TIL10 | TIL08 | BKN022 | TDD39103 | 30368 | 40744 | -0.1459107 | 40.32787 | 0.00361811 |
| TIL11 | TIL08 | BKN022 | TDD39103 | 32367 | 42820 | -0.1390267 | 37.50644 | 0.003706742 |
| TIL12 | TIL08 | BKN022 | TDD39103 | 28874 | 33070 | -0.0677386 | 17.19538 | 0.00393935 |
| TIL14-TIP498 | TIL08 | BKN022 | TDD39103 | 31682 | 42851 | -0.1498531 | 42.22892 | 0.003548589 |
| TIL15 | TIL08 | BKN022 | TDD39103 | 32230 | 35564 | -0.04917839 | 13.5574 | 0.00362742 |
| TIL16 | TIL08 | BKN022 | TDD39103 | 31647 | 33781 | -0.03261601 | 7.888716 | 0.004134514 |
| TIL17 | TIL08 | BKN022 | TDD39103 | 29262 | 41352 | -0.1712125 | 40.29914 | 0.00424854 |
| TIL01 | TIL25-TIP489 | BKN022 | TDD39103 | 31539 | 41245 | -0.1333535 | 35.76747 | 0.003728345 |

*Continuation of Table S7*

| *parviglumis* | *mexicana* | Test | Outgroup | ABBA sites | BABA sites | D-raw | Z-score | SD-D statistic |
|---|---|---|---|---|---|---|---|---|
| TIL01-JD | TIL25-TIP489 | BKN022 | TDD39103 | 32273 | 42386 | -0.1354559 | 33.10467 | 0.004091745 |
| TIL02 | TIL25-TIP489 | BKN022 | TDD39103 | 27264 | 38059 | -0.1652557 | 41.73333 | 0.003959802 |
| TIL03 | TIL25-TIP489 | BKN022 | TDD39103 | 31263 | 48379 | -0.2149117 | 66.92783 | 0.003211096 |
| TIL04-TIP454 | TIL25-TIP489 | BKN022 | TDD39103 | 30817 | 42039 | -0.1540299 | 40.30662 | 0.003821453 |
| TIL05 | TIL25-TIP489 | BKN022 | TDD39103 | 31628 | 42220 | -0.1434297 | 37.452 | 0.003829696 |
| TIL06-TIP260 | TIL25-TIP489 | BKN022 | TDD39103 | 31713 | 36888 | -0.07543622 | 20.08414 | 0.00375601 |
| TIL06-TIP496 | TIL25-TIP489 | BKN022 | TDD39103 | 31164 | 36827 | -0.08329044 | 23.24494 | 0.003583165 |
| TIL07 | TIL25-TIP489 | BKN022 | TDD39103 | 32074 | 43314 | -0.1490953 | 37.43665 | 0.003982604 |
| TIL09 | TIL25-TIP489 | BKN022 | TDD39103 | 32332 | 43913 | -0.1518919 | 40.50072 | 0.003750351 |
| TIL10 | TIL25-TIP489 | BKN022 | TDD39103 | 31608 | 43124 | -0.1540973 | 41.92581 | 0.003675476 |
| TIL11 | TIL25-TIP489 | BKN022 | TDD39103 | 32229 | 43806 | -0.1522588 | 48.90107 | 0.003113609 |
| TIL12 | TIL25-TIP489 | BKN022 | TDD39103 | 29015 | 34357 | -0.0842959 | 23.12754 | 0.003644828 |
| TIL14-TIP498 | TIL25-TIP489 | BKN022 | TDD39103 | 31888 | 44042 | -0.1600685 | 42.82319 | 0.003737893 |
| TIL15 | TIL25-TIP489 | BKN022 | TDD39103 | 31876 | 36171 | -0.06311814 | 16.51133 | 0.003822717 |
| TIL16 | TIL25-TIP489 | BKN022 | TDD39103 | 31421 | 34445 | -0.0459114 | 13.28367 | 0.003456228 |
| TIL17 | TIL25-TIP489 | BKN022 | TDD39103 | 30352 | 43606 | -0.1792098 | 48.05557 | 0.00372922 |

**Table S8.** D-statistic values for each pairwise combination of (TEST, *parviglumis*, *mexicana*, outgroup).

| *parviglumis* | *mexicana* | Test | Outgroup | ABBA sites | BABA sites | D-raw | Z-score | SD-D statistic |
|---|---|---|---|---|---|---|---|---|
| TIL01 | TIL08 | Par_N16 | TDD39103 | 46436 | 25347 | 0.2937882 | 77.94676 | 0.003769088 |
| TIL01-JD | TIL08 | Par_N16 | TDD39103 | 48633 | 26111 | 0.3013218 | 85.54295 | 0.003522463 |
| TIL02 | TIL08 | Par_N16 | TDD39103 | 40056 | 19045 | 0.3555101 | 90.34326 | 0.003935103 |
| TIL03 | TIL08 | Par_N16 | TDD39103 | 45948 | 24486 | 0.3047108 | 92.17341 | 0.003305843 |
| TIL04-TIP454 | TIL08 | Par_N16 | TDD39103 | 52597 | 22866 | 0.3939812 | 101.0925 | 0.003897235 |
| TIL05 | TIL08 | Par_N16 | TDD39103 | 55415 | 25085 | 0.3767702 | 117.8533 | 0.003196941 |
| TIL06-TIP260 | TIL08 | Par_N16 | TDD39103 | 55062 | 25174 | 0.3725011 | 105.0468 | 0.003546048 |
| TIL06-TIP496 | TIL08 | Par_N16 | TDD39103 | 54351 | 24770 | 0.3738704 | 112.6592 | 0.003318596 |
| TIL07 | TIL08 | Par_N16 | TDD39103 | 48294 | 23966 | 0.3366731 | 109.4535 | 0.003075945 |
| TIL09 | TIL08 | Par_N16 | TDD39103 | 49094 | 24969 | 0.3257362 | 93.87043 | 0.003470062 |
| TIL10 | TIL08 | Par_N16 | TDD39103 | 55442 | 23596 | 0.4029201 | 127.4567 | 0.003161231 |
| TIL11 | TIL08 | Par_N16 | TDD39103 | 47251 | 25461 | 0.2996754 | 101.1621 | 0.002962328 |
| TIL12 | TIL08 | Par_N16 | TDD39103 | 56984 | 22925 | 0.4262223 | 131.8929 | 0.00323158 |
| TIL14-TIP498 | TIL08 | Par_N16 | TDD39103 | 47398 | 25207 | 0.3056401 | 90.6943 | 0.003370003 |
| TIL15 | TIL08 | Par_N16 | TDD39103 | 59440 | 26198 | 0.3881688 | 122.9034 | 0.003158324 |
| TIL16 | TIL08 | Par_N16 | TDD39103 | 51842 | 25179 | 0.3461783 | 97.18702 | 0.003561981 |
| TIL17 | TIL08 | Par_N16 | TDD39103 | 48957 | 22501 | 0.3702315 | 109.4335 | 0.003383164 |
| TIL01 | TIL25-TIP489 | Par_N16 | TDD39103 | 55981 | 23853 | 0.4024351 | 113.9415 | 0.003531944 |
| TIL01-JD | TIL25-TIP489 | Par_N16 | TDD39103 | 59555 | 24571 | 0.4158524 | 134.0795 | 0.003101537 |
| TIL02 | TIL25-TIP489 | Par_N16 | TDD39103 | 42937 | 19117 | 0.3838592 | 119.5291 | 0.003211429 |
| TIL03 | TIL25-TIP489 | Par_N16 | TDD39103 | 53953 | 23846 | 0.3869844 | 125.2472 | 0.003089765 |
| TIL04-TIP454 | TIL25-TIP489 | Par_N16 | TDD39103 | 57281 | 22989 | 0.4272082 | 126.2158 | 0.003384744 |

*Continuation of Table S8*

| *parviglumis* | *mexicana* | Test | Outgroup | ABBA sites | BABA sites | D-raw | Z-score | SD-D statistic |
|---|---|---|---|---|---|---|---|---|
| TIL05 | TIL25-TIP489 | Par_N16 | TDD39103 | 63309 | 23990 | 0.4503946 | 140.4084 | 0.003207748 |
| TIL06-TIP260 | TIL25-TIP489 | Par_N16 | TDD39103 | 63523 | 24110 | 0.4497507 | 162.9832 | 0.002759491 |
| TIL06-TIP496 | TIL25-TIP489 | Par_N16 | TDD39103 | 62625 | 23838 | 0.4485965 | 127.825 | 0.003509459 |
| TIL07 | TIL25-TIP489 | Par_N16 | TDD39103 | 49692 | 24345 | 0.3423558 | 98.34875 | 0.003481039 |
| TIL09 | TIL25-TIP489 | Par_N16 | TDD39103 | 55244 | 24538 | 0.3848738 | 129.2032 | 0.002978825 |
| TIL10 | TIL25-TIP489 | Par_N16 | TDD39103 | 59278 | 23906 | 0.425226 | 149.7145 | 0.002840245 |
| TIL11 | TIL25-TIP489 | Par_N16 | TDD39103 | 61998 | 24281 | 0.4371516 | 151.2681 | 0.002889912 |
| TIL12 | TIL25-TIP489 | Par_N16 | TDD39103 | 61461 | 21730 | 0.4775877 | 163.653 | 0.002918295 |
| TIL14-TIP498 | TIL25-TIP489 | Par_N16 | TDD39103 | 61538 | 24261 | 0.4344689 | 143.4789 | 0.003028104 |
| TIL15 | TIL25-TIP489 | Par_N16 | TDD39103 | 71366 | 24530 | 0.4884041 | 179.9939 | 0.002713448 |
| TIL16 | TIL25-TIP489 | Par_N16 | TDD39103 | 62059 | 23666 | 0.4478624 | 152.5426 | 0.002935982 |
| TIL17 | TIL25-TIP489 | Par_N16 | TDD39103 | 49765 | 22668 | 0.3740974 | 113.5682 | 0.003294034 |
| TIL01 | TIL08 | PT2233 | TDD39103 | 7896 | 14543 | -0.2962253 | 45.90482 | 0.006453033 |
| TIL01-JD | TIL08 | PT2233 | TDD39103 | 8056 | 14743 | -0.2933023 | 44.78754 | 0.006548748 |
| TIL02 | TIL08 | PT2233 | TDD39103 | 7428 | 13367 | -0.2855975 | 44.80682 | 0.006373973 |
| TIL03 | TIL08 | PT2233 | TDD39103 | 7906 | 14856 | -0.3053335 | 48.57318 | 0.00628605 |
| TIL04-TIP454 | TIL08 | PT2233 | TDD39103 | 8308 | 13793 | -0.2481788 | 39.69596 | 0.006251992 |
| TIL05 | TIL08 | PT2233 | TDD39103 | 8362 | 14301 | -0.2620571 | 45.9842 | 0.00569885 |
| TIL06-TIP260 | TIL08 | PT2233 | TDD39103 | 8239 | 13808 | -0.2525967 | 40.15552 | 0.006290461 |
| TIL06-TIP496 | TIL08 | PT2233 | TDD39103 | 8129 | 13671 | -0.2542202 | 42.1049 | 0.006037782 |
| TIL07 | TIL08 | PT2233 | TDD39103 | 7978 | 14296 | -0.2836491 | 43.32269 | 0.006547357 |
| TIL09 | TIL08 | PT2233 | TDD39103 | 8108 | 14542 | -0.2840618 | 43.88435 | 0.006472964 |
| TIL10 | TIL08 | PT2233 | TDD39103 | 8412 | 14086 | -0.2522002 | 40.22137 | 0.006270303 |
| TIL11 | TIL08 | PT2233 | TDD39103 | 8074 | 14624 | -0.2885717 | 50.40609 | 0.005724937 |
| TIL12 | TIL08 | PT2233 | TDD39103 | 8116 | 13298 | -0.2419912 | 32.96327 | 0.007341239 |

*Continuation of Table S8*

| *parviglumis* | *mexicana* | Test | Outgroup | ABBA sites | BABA sites | D-raw | Z-score | SD-D statistic |
|---|---|---|---|---|---|---|---|---|
| TIL14-TIP498 | TIL08 | PT2233 | TDD39103 | 8187 | 14644 | -0.2828172 | 42.11459 | 0.006715421 |
| TIL15 | TIL08 | PT2233 | TDD39103 | 8403 | 13621 | -0.2369234 | 34.78425 | 0.006811226 |
| TIL16 | TIL08 | PT2233 | TDD39103 | 7970 | 13779 | -0.2670927 | 42.14891 | 0.006336884 |
| TIL17 | TIL08 | PT2233 | TDD39103 | 8111 | 13972 | -0.2654078 | 42.16933 | 0.006293858 |
| TIL01 | TIL25-TIP489 | PT2233 | TDD39103 | 7880 | 16455 | -0.3523731 | 55.41818 | 0.006358439 |
| TIL01-JD | TIL25-TIP489 | PT2233 | TDD39103 | 8157 | 16619 | -0.3415402 | 58.90813 | 0.005797845 |
| TIL02 | TIL25-TIP489 | PT2233 | TDD39103 | 7335 | 15414 | -0.3551365 | 63.09221 | 0.005628848 |
| TIL03 | TIL25-TIP489 | PT2233 | TDD39103 | 7896 | 16630 | -0.3561119 | 57.68866 | 0.006172996 |
| TIL04-TIP454 | TIL25-TIP489 | PT2233 | TDD39103 | 8119 | 15987 | -0.3263918 | 53.88356 | 0.006057354 |
| TIL05 | TIL25-TIP489 | PT2233 | TDD39103 | 8325 | 16203 | -0.321184 | 54.12132 | 0.005934518 |
| TIL06-TIP260 | TIL25-TIP489 | PT2233 | TDD39103 | 8142 | 16009 | -0.3257422 | 52.92274 | 0.006155052 |
| TIL06-TIP496 | TIL25-TIP489 | PT2233 | TDD39103 | 8042 | 15889 | -0.327901 | 59.08939 | 0.005549237 |
| TIL07 | TIL25-TIP489 | PT2233 | TDD39103 | 7789 | 16566 | -0.3603777 | 66.78656 | 0.005395962 |
| TIL09 | TIL25-TIP489 | PT2233 | TDD39103 | 8163 | 16443 | -0.3365033 | 52.64749 | 0.00639163 |
| TIL10 | TIL25-TIP489 | PT2233 | TDD39103 | 8250 | 16227 | -0.3258978 | 52.39601 | 0.006219897 |
| TIL11 | TIL25-TIP489 | PT2233 | TDD39103 | 8093 | 16457 | -0.3406925 | 63.15931 | 0.005394177 |
| TIL12 | TIL25-TIP489 | PT2233 | TDD39103 | 8145 | 15222 | -0.302863 | 41.76151 | 0.007252205 |
| TIL14-TIP498 | TIL25-TIP489 | PT2233 | TDD39103 | 8136 | 16565 | -0.3412412 | 59.53785 | 0.005731501 |
| TIL15 | TIL25-TIP489 | PT2233 | TDD39103 | 8543 | 15617 | -0.292798 | 45.54131 | 0.006429284 |
| TIL16 | TIL25-TIP489 | PT2233 | TDD39103 | 8101 | 15589 | -0.3160827 | 48.21893 | 0.006555158 |
| TIL17 | TIL25-TIP489 | PT2233 | TDD39103 | 7938 | 16221 | -0.3428536 | 56.58369 | 0.00605923 |
| TIL01 | TIL08 | BKN022 | TDD39103 | 41629 | 37987 | 0.04574457 | 12.65302 | 0.003615309 |
| TIL01-JD | TIL08 | BKN022 | TDD39103 | 43514 | 38843 | 0.05671649 | 15.29806 | 0.003707431 |
| TIL02 | TIL08 | BKN022 | TDD39103 | 35853 | 31036 | 0.07201483 | 17.53659 | 0.004106546 |
| TIL03 | TIL08 | BKN022 | TDD39103 | 40187 | 36493 | 0.04817423 | 13.38069 | 0.00360028 |

*Continuation of Table S8*

| *parviglumis* | *mexicana* | Test | Outgroup | ABBA sites | BABA sites | D-raw | Z-score | SD-D statistic |
|---|---|---|---|---|---|---|---|---|
| TIL04-TIP454 | TIL08 | BKN022 | TDD39103 | 47384 | 34718 | 0.1542715 | 45.83063 | 0.003366123 |
| TIL05 | TIL08 | BKN022 | TDD39103 | 49721 | 37040 | 0.1461601 | 38.08849 | 0.003837383 |
| TIL06-TIP260 | TIL08 | BKN022 | TDD39103 | 49880 | 37042 | 0.1476956 | 38.14916 | 0.003871531 |
| TIL06-TIP496 | TIL08 | BKN022 | TDD39103 | 48975 | 36374 | 0.1476409 | 44.04696 | 0.003351896 |
| TIL07 | TIL08 | BKN022 | TDD39103 | 42939 | 35896 | 0.08933849 | 21.54374 | 0.004146842 |
| TIL09 | TIL08 | BKN022 | TDD39103 | 44352 | 38052 | 0.0764526 | 23.06499 | 0.00331466 |
| TIL10 | TIL08 | BKN022 | TDD39103 | 50577 | 36290 | 0.1644698 | 51.90085 | 0.003168923 |
| TIL11 | TIL08 | BKN022 | TDD39103 | 42461 | 38432 | 0.04980653 | 13.04113 | 0.003819189 |
| TIL12 | TIL08 | BKN022 | TDD39103 | 51179 | 34016 | 0.2014555 | 68.88469 | 0.002924532 |
| TIL14-TIP498 | TIL08 | BKN022 | TDD39103 | 41964 | 37485 | 0.05637579 | 15.1802 | 0.003713771 |
| TIL15 | TIL08 | BKN022 | TDD39103 | 53848 | 38098 | 0.1712962 | 49.26255 | 0.003477209 |
| TIL16 | TIL08 | BKN022 | TDD39103 | 46773 | 37101 | 0.1153158 | 30.17569 | 0.003821481 |
| TIL17 | TIL08 | BKN022 | TDD39103 | 43742 | 34519 | 0.1178492 | 34.78665 | 0.003387773 |
| TIL01 | TIL25-TIP489 | BKN022 | TDD39103 | 50367 | 38756 | 0.1302806 | 35.29869 | 0.003690806 |
| TIL01-JD | TIL25-TIP489 | BKN022 | TDD39103 | 53531 | 39569 | 0.1499678 | 45.81329 | 0.003273456 |
| TIL02 | TIL25-TIP489 | BKN022 | TDD39103 | 38043 | 33221 | 0.0676639 | 17.78616 | 0.003804299 |
| TIL03 | TIL25-TIP489 | BKN022 | TDD39103 | 47352 | 38276 | 0.1059934 | 32.3813 | 0.003273289 |
| TIL04-TIP454 | TIL25-TIP489 | BKN022 | TDD39103 | 51326 | 37335 | 0.1578033 | 48.34967 | 0.003263793 |
| TIL05 | TIL25-TIP489 | BKN022 | TDD39103 | 57157 | 38618 | 0.1935683 | 62.48952 | 0.003097612 |
| TIL06-TIP260 | TIL25-TIP489 | BKN022 | TDD39103 | 57557 | 38492 | 0.1984924 | 62.85717 | 0.003157833 |
| TIL06-TIP496 | TIL25-TIP489 | BKN022 | TDD39103 | 56451 | 37887 | 0.1967818 | 56.81167 | 0.003463757 |
| TIL07 | TIL25-TIP489 | BKN022 | TDD39103 | 43405 | 38800 | 0.05601849 | 15.36165 | 0.003646645 |
| TIL09 | TIL25-TIP489 | BKN022 | TDD39103 | 49102 | 39646 | 0.1065489 | 27.86222 | 0.003824135 |
| TIL10 | TIL25-TIP489 | BKN022 | TDD39103 | 53024 | 38667 | 0.1565803 | 47.16362 | 0.003319937 |

*Continuation of Table S8*

| *parviglumis* | *mexicana* | Test | Outgroup | ABBA sites | BABA sites | D-raw | Z-score | SD-D statistic |
|---|---|---|---|---|---|---|---|---|
| TIL11 | TIL25-TIP489 | BKN022 | TDD39103 | 56416 | 39612 | 0.1749906 | 57.75036 | 0.003030122 |
| TIL12 | TIL25-TIP489 | BKN022 | TDD39103 | 55280 | 35269 | 0.2209964 | 71.55078 | 0.003088665 |
| TIL14-TIP498 | TIL25-TIP489 | BKN022 | TDD39103 | 55308 | 38963 | 0.1733831 | 61.3781 | 0.002824837 |
| TIL15 | TIL25-TIP489 | BKN022 | TDD39103 | 65205 | 39132 | 0.2498922 | 90.4434 | 0.002762968 |
| TIL16 | TIL25-TIP489 | BKN022 | TDD39103 | 56465 | 38222 | 0.1926664 | 60.90295 | 0.003163498 |
| TIL17 | TIL25-TIP489 | BKN022 | TDD39103 | 43630 | 36891 | 0.08369245 | 24.88248 | 0.003363509 |

**Table S9.** D-statistic values for each pairwise combination of (TEST, *parviglumis*, *mexicana*, outgroup) including only transversions.

| *parviglumis* | *mexicana* | Test | Outgroup | ABBA sites | BABA sites | D-raw | Z-score | SD-D statistic |
|---|---|---|---|---|---|---|---|---|
| TIL01 | TIL08 | Par_N16 | TDD39103 | 38397 | 23899 | 0.2327276 | 65.33946 | 0.003561824 |
| TIL01-JD | TIL08 | Par_N16 | TDD39103 | 40114 | 24663 | 0.238526 | 64.24741 | 0.003712617 |
| TIL02 | TIL08 | Par_N16 | TDD39103 | 33170 | 18079 | 0.2944643 | 66.75864 | 0.004410879 |
| TIL03 | TIL08 | Par_N16 | TDD39103 | 37932 | 23000 | 0.2450601 | 65.03308 | 0.003768237 |
| TIL04-TIP454 | TIL08 | Par_N16 | TDD39103 | 43253 | 21569 | 0.3345161 | 109.8866 | 0.003044194 |
| TIL05 | TIL08 | Par_N16 | TDD39103 | 45152 | 23650 | 0.31252 | 83.4583 | 0.003744624 |
| TIL06-TIP260 | TIL08 | Par_N16 | TDD39103 | 45373 | 23933 | 0.3093527 | 81.11714 | 0.003813654 |
| TIL06-TIP496 | TIL08 | Par_N16 | TDD39103 | 44753 | 23427 | 0.3127897 | 87.81918 | 0.003561747 |
| TIL07 | TIL08 | Par_N16 | TDD39103 | 40220 | 22781 | 0.2768051 | 77.06923 | 0.003591642 |
| TIL09 | TIL08 | Par_N16 | TDD39103 | 39971 | 23646 | 0.2566138 | 68.05175 | 0.003770863 |
| TIL10 | TIL08 | Par_N16 | TDD39103 | 45003 | 22283 | 0.3376631 | 85.83785 | 0.003933732 |
| TIL11 | TIL08 | Par_N16 | TDD39103 | 39158 | 23950 | 0.2409837 | 59.43876 | 0.00405432 |
| TIL12 | TIL08 | Par_N16 | TDD39103 | 47117 | 21798 | 0.3673946 | 101.9182 | 0.003604799 |
| TIL14-TIP498 | TIL08 | Par_N16 | TDD39103 | 39250 | 23764 | 0.2457549 | 57.05173 | 0.004307581 |
| TIL15 | TIL08 | Par_N16 | TDD39103 | 48746 | 24928 | 0.3232891 | 104.5397 | 0.003092499 |
| TIL16 | TIL08 | Par_N16 | TDD39103 | 43274 | 23929 | 0.2878592 | 76.29583 | 0.003772935 |
| TIL17 | TIL08 | Par_N16 | TDD39103 | 40533 | 21301 | 0.3110263 | 79.72286 | 0.003901344 |
| TIL01 | TIL25-TIP489 | Par_N16 | TDD39103 | 45853 | 22070 | 0.3501465 | 86.58404 | 0.004044007 |
| TIL01-JD | TIL25-TIP489 | Par_N16 | TDD39103 | 48427 | 22788 | 0.3600225 | 99.98153 | 0.00360089 |
| TIL02 | TIL25-TIP489 | Par_N16 | TDD39103 | 34962 | 17717 | 0.32736 | 79.99527 | 0.004092242 |
| TIL03 | TIL25-TIP489 | Par_N16 | TDD39103 | 42769 | 22054 | 0.3195625 | 88.06544 | 0.003628694 |
| TIL04-TIP454 | TIL25-TIP489 | Par_N16 | TDD39103 | 45883 | 21390 | 0.3640837 | 105.6736 | 0.003445362 |

*Continuation of Table S9*

| *parviglumis* | *mexicana* | Test | Outgroup | ABBA sites | BABA sites | D-raw | Z-score | SD-D statistic |
|---|---|---|---|---|---|---|---|---|
| TIL05 | TIL25-TIP489 | Par_N16 | TDD39103 | 50889 | 22242 | 0.3917217 | 106.6289 | 0.003673692 |
| TIL06-TIP260 | TIL25-TIP489 | Par_N16 | TDD39103 | 52050 | 22530 | 0.3958166 | 121.0721 | 0.003269262 |
| TIL06-TIP496 | TIL25-TIP489 | Par_N16 | TDD39103 | 51031 | 22126 | 0.3951091 | 111.1652 | 0.003554253 |
| TIL07 | TIL25-TIP489 | Par_N16 | TDD39103 | 40777 | 22702 | 0.2847398 | 68.64007 | 0.004148304 |
| TIL09 | TIL25-TIP489 | Par_N16 | TDD39103 | 44562 | 22931 | 0.3204925 | 104.9692 | 0.003053206 |
| TIL10 | TIL25-TIP489 | Par_N16 | TDD39103 | 46595 | 22179 | 0.3550179 | 92.93927 | 0.003819891 |
| TIL11 | TIL25-TIP489 | Par_N16 | TDD39103 | 49837 | 22477 | 0.37835 | 117.5801 | 0.003217806 |
| TIL12 | TIL25-TIP489 | Par_N16 | TDD39103 | 51345 | 20299 | 0.4333371 | 129.0593 | 0.003357659 |
| TIL14-TIP498 | TIL25-TIP489 | Par_N16 | TDD39103 | 48880 | 22477 | 0.3700128 | 105.694 | 0.003500793 |
| TIL15 | TIL25-TIP489 | Par_N16 | TDD39103 | 58088 | 22965 | 0.4333337 | 139.3251 | 0.003110235 |
| TIL16 | TIL25-TIP489 | Par_N16 | TDD39103 | 51493 | 22049 | 0.4003699 | 133.5687 | 0.002997482 |
| TIL17 | TIL25-TIP489 | Par_N16 | TDD39103 | 40591 | 21044 | 0.3171412 | 78.92576 | 0.004018222 |
| TIL01 | TIL08 | PT2233 | TDD39103 | 37169 | 30154 | 0.1041992 | 28.29494 | 0.003682608 |
| TIL01-JD | TIL08 | PT2233 | TDD39103 | 38783 | 31004 | 0.1114678 | 36.13437 | 0.003084812 |
| TIL02 | TIL08 | PT2233 | TDD39103 | 32414 | 24276 | 0.1435527 | 34.74403 | 0.004131722 |
| TIL03 | TIL08 | PT2233 | TDD39103 | 36766 | 29429 | 0.1108392 | 26.47428 | 0.004186674 |
| TIL04-TIP454 | TIL08 | PT2233 | TDD39103 | 42033 | 27659 | 0.2062504 | 57.98201 | 0.003557144 |
| TIL05 | TIL08 | PT2233 | TDD39103 | 43765 | 29717 | 0.1911761 | 52.29399 | 0.003655794 |
| TIL06-TIP260 | TIL08 | PT2233 | TDD39103 | 43877 | 29740 | 0.1920344 | 57.43864 | 0.003343297 |
| TIL06-TIP496 | TIL08 | PT2233 | TDD39103 | 43248 | 29135 | 0.1949767 | 55.82905 | 0.003492388 |
| TIL07 | TIL08 | PT2233 | TDD39103 | 38960 | 29045 | 0.1457981 | 39.54577 | 0.003686819 |
| TIL09 | TIL08 | PT2233 | TDD39103 | 38672 | 29943 | 0.1272171 | 35.28027 | 0.003605898 |
| TIL10 | TIL08 | PT2233 | TDD39103 | 43697 | 28477 | 0.2108793 | 57.53387 | 0.003665306 |
| TIL11 | TIL08 | PT2233 | TDD39103 | 38029 | 30355 | 0.1122192 | 29.96077 | 0.003745539 |
| TIL12 | TIL08 | PT2233 | TDD39103 | 45493 | 27349 | 0.2490871 | 68.15686 | 0.003654615 |

*Continuation of Table S9*

| *parviglumis* | *mexicana* | Test | Outgroup | ABBA sites | BABA sites | D-raw | Z-score | SD-D statistic |
|---|---|---|---|---|---|---|---|---|
| TIL14-TIP498 | TIL08 | PT2233 | TDD39103 | 38177 | 30189 | 0.1168417 | 30.41161 | 0.00384201 |
| TIL15 | TIL08 | PT2233 | TDD39103 | 46763 | 30413 | 0.2118534 | 58.85992 | 0.003599281 |
| TIL16 | TIL08 | PT2233 | TDD39103 | 41724 | 29699 | 0.1683631 | 42.48423 | 0.003962956 |
| TIL17 | TIL08 | PT2233 | TDD39103 | 39411 | 27538 | 0.1773439 | 43.77723 | 0.004051055 |
| TIL01 | TIL25-TIP489 | PT2233 | TDD39103 | 44197 | 30159 | 0.1887944 | 52.99782 | 0.003562306 |
| TIL01-JD | TIL25-TIP489 | PT2233 | TDD39103 | 46619 | 30866 | 0.2033039 | 50.7704 | 0.004004378 |
| TIL02 | TIL25-TIP489 | PT2233 | TDD39103 | 33730 | 25818 | 0.1328676 | 32.91049 | 0.004037242 |
| TIL03 | TIL25-TIP489 | PT2233 | TDD39103 | 41092 | 30258 | 0.151843 | 39.96822 | 0.003799094 |
| TIL04-TIP454 | TIL25-TIP489 | PT2233 | TDD39103 | 44187 | 29340 | 0.2019258 | 55.87255 | 0.003614043 |
| TIL05 | TIL25-TIP489 | PT2233 | TDD39103 | 49013 | 30097 | 0.2391101 | 66.2672 | 0.003608272 |
| TIL06-TIP260 | TIL25-TIP489 | PT2233 | TDD39103 | 50233 | 30285 | 0.2477458 | 78.28821 | 0.003164536 |
| TIL06-TIP496 | TIL25-TIP489 | PT2233 | TDD39103 | 49226 | 29806 | 0.2457233 | 68.4503 | 0.003589805 |
| TIL07 | TIL25-TIP489 | PT2233 | TDD39103 | 39045 | 30910 | 0.116289 | 28.55518 | 0.004072432 |
| TIL09 | TIL25-TIP489 | PT2233 | TDD39103 | 42740 | 30948 | 0.1600261 | 43.76523 | 0.003656465 |
| TIL10 | TIL25-TIP489 | PT2233 | TDD39103 | 44668 | 30110 | 0.1946829 | 49.02424 | 0.003971156 |
| TIL11 | TIL25-TIP489 | PT2233 | TDD39103 | 48126 | 30527 | 0.223755 | 58.45579 | 0.003827764 |
| TIL12 | TIL25-TIP489 | PT2233 | TDD39103 | 49410 | 27785 | 0.2801347 | 74.37125 | 0.003766707 |
| TIL14-TIP498 | TIL25-TIP489 | PT2233 | TDD39103 | 47331 | 30681 | 0.2134287 | 52.79837 | 0.004042335 |
| TIL15 | TIL25-TIP489 | PT2233 | TDD39103 | 55663 | 30264 | 0.2955881 | 93.55037 | 0.003159668 |
| TIL16 | TIL25-TIP489 | PT2233 | TDD39103 | 49496 | 29586 | 0.251764 | 82.57528 | 0.003048903 |
| TIL17 | TIL25-TIP489 | PT2233 | TDD39103 | 39068 | 29294 | 0.1429742 | 39.4705 | 0.003622304 |
| TIL01 | TIL08 | BKN022 | TDD39103 | 34404 | 31935 | 0.03721793 | 9.332679 | 0.003987914 |
| TIL01-JD | TIL08 | BKN022 | TDD39103 | 35843 | 32703 | 0.04580865 | 10.41325 | 0.004399075 |
| TIL02 | TIL08 | BKN022 | TDD39103 | 30179 | 26125 | 0.07200199 | 17.24475 | 0.004175299 |
| TIL03 | TIL08 | BKN022 | TDD39103 | 33339 | 30644 | 0.04212056 | 10.76355 | 0.003913259 |

*Continuation of Table S9*

| *parviglumis* | *mexicana* | Test | Outgroup | ABBA sites | BABA sites | D-raw | Z-score | SD-D statistic |
|---|---|---|---|---|---|---|---|---|
| TIL04-TIP454 | TIL08 | BKN022 | TDD39103 | 39277 | 29362 | 0.1444514 | 36.67931 | 0.003938226 |
| TIL05 | TIL08 | BKN022 | TDD39103 | 40596 | 31263 | 0.1298793 | 28.35329 | 0.004580751 |
| TIL06-TIP260 | TIL08 | BKN022 | TDD39103 | 41164 | 31507 | 0.1328866 | 42.02312 | 0.003162226 |
| TIL06-TIP496 | TIL08 | BKN022 | TDD39103 | 40443 | 30840 | 0.1347166 | 36.61385 | 0.003679388 |
| TIL07 | TIL08 | BKN022 | TDD39103 | 35992 | 30506 | 0.08249872 | 23.88856 | 0.003453483 |
| TIL09 | TIL08 | BKN022 | TDD39103 | 36138 | 32023 | 0.06037177 | 15.00756 | 0.004022757 |
| TIL10 | TIL08 | BKN022 | TDD39103 | 40954 | 30368 | 0.1484255 | 41.23392 | 0.003599596 |
| TIL11 | TIL08 | BKN022 | TDD39103 | 35384 | 32367 | 0.04453071 | 12.17695 | 0.003656966 |
| TIL12 | TIL08 | BKN022 | TDD39103 | 42542 | 28874 | 0.1913857 | 48.57234 | 0.00394022 |
| TIL14-TIP498 | TIL08 | BKN022 | TDD39103 | 35026 | 31682 | 0.05012892 | 15.36072 | 0.003263448 |
| TIL15 | TIL08 | BKN022 | TDD39103 | 43927 | 32230 | 0.1535906 | 39.2013 | 0.003917998 |
| TIL16 | TIL08 | BKN022 | TDD39103 | 39286 | 31647 | 0.1076932 | 28.98477 | 0.00371551 |
| TIL17 | TIL08 | BKN022 | TDD39103 | 36614 | 29262 | 0.1116036 | 29.15789 | 0.003827561 |
| TIL01 | TIL25-TIP489 | BKN022 | TDD39103 | 41412 | 31539 | 0.1353374 | 36.06972 | 0.003752106 |
| TIL01-JD | TIL25-TIP489 | BKN022 | TDD39103 | 43658 | 32273 | 0.1499388 | 43.15498 | 0.003474426 |
| TIL02 | TIL25-TIP489 | BKN022 | TDD39103 | 31474 | 27264 | 0.07167421 | 20.76299 | 0.003452018 |
| TIL03 | TIL25-TIP489 | BKN022 | TDD39103 | 37705 | 31263 | 0.09340564 | 30.19402 | 0.003093515 |
| TIL04-TIP454 | TIL25-TIP489 | BKN022 | TDD39103 | 41428 | 30817 | 0.1468752 | 40.14434 | 0.003658678 |
| TIL05 | TIL25-TIP489 | BKN022 | TDD39103 | 46132 | 31628 | 0.1865226 | 50.72378 | 0.003677223 |
| TIL06-TIP260 | TIL25-TIP489 | BKN022 | TDD39103 | 47337 | 31713 | 0.1976471 | 59.89253 | 0.003300028 |
| TIL06-TIP496 | TIL25-TIP489 | BKN022 | TDD39103 | 46268 | 31164 | 0.1950615 | 56.49287 | 0.003452851 |
| TIL07 | TIL25-TIP489 | BKN022 | TDD39103 | 36025 | 32074 | 0.05801847 | 15.37299 | 0.003774052 |
| TIL09 | TIL25-TIP489 | BKN022 | TDD39103 | 39661 | 32332 | 0.1018016 | 24.38927 | 0.004174031 |
| TIL10 | TIL25-TIP489 | BKN022 | TDD39103 | 41709 | 31608 | 0.1377716 | 38.13418 | 0.003612811 |

*Continuation of Table S9*

| *parviglumis* | *mexicana* | Test | Outgroup | ABBA sites | BABA sites | D-raw | Z-score | SD-D statistic |
|---|---|---|---|---|---|---|---|---|
| TIL11 | TIL25-TIP489 | BKN022 | TDD39103 | 45468 | 32229 | 0.1703927 | 46.08421 | 0.00369742 |
| TIL12 | TIL25-TIP489 | BKN022 | TDD39103 | 46583 | 29015 | 0.2323871 | 68.24823 | 0.003405027 |
| TIL14-TIP498 | TIL25-TIP489 | BKN022 | TDD39103 | 44115 | 31888 | 0.1608752 | 45.01638 | 0.003573704 |
| TIL15 | TIL25-TIP489 | BKN022 | TDD39103 | 52836 | 31876 | 0.2474266 | 74.09266 | 0.003339421 |
| TIL16 | TIL25-TIP489 | BKN022 | TDD39103 | 47164 | 31421 | 0.2003309 | 56.81827 | 0.003525818 |
| TIL17 | TIL25-TIP489 | BKN022 | TDD39103 | 36040 | 30352 | 0.08567297 | 21.77772 | 0.003933973 |

**Table S10.** Allele frequency of highland and lowland Mesoamerican landraces at sites intersected with Par_N16 from 668 adaptive SNPs.

| Chromosome | Position | Par_N16 genotype | Allele frequency highlands | Allele frequency lowlands |
|---|---|---|---|---|
| 4 | 178417584 | TT | 0.3823529412 | 1 |
| 4 | 170368667 | CC | 0.5 | 1 |
| 8 | 145009951 | TT | 1 | 0.3181818182 |
| 1 | 149125711 | TT | 0.7916666667 | 0.1 |
| 8 | 67810183 | TT | 0.7916666667 | 0.5909090909 |
| 8 | 19685236 | GG | 0.5 | 0.9 |
| 4 | 180529091 | AA | 0.3333333333 | 1 |
| 3 | 154634811 | AA | 0.3333333333 | 0.8611111111 |
| 5 | 76968662 | CC | 0.15 | 0.8846153846 |
| 8 | 161076561 | CC | 0.2142857143 | 0.85 |
| 2 | 234268487 | GG | 0.4736842105 | 0.9736842105 |
| 8 | 2517850 | GG | 0.9583333333 | 0.4444444444 |
| 4 | 180529051 | AA | 0.3333333333 | 1 |
| 10 | 8455981 | CC | 0.28125 | 0.9210526316 |
| 3 | 191200343 | TT | 0.4166666667 | 0.9210526316 |
| 7 | 114903332 | CC | 0.9583333333 | 0.3846153846 |
| 2 | 74938106 | CC | 0.1052631579 | 0.7142857143 |
| 4 | 169577208 | GG | 0.2083333333 | 0.8846153846 |
| 4 | 177090360 | GG | 0.5 | 1 |
| 8 | 111325446 | CC | 0.1875 | 0.6666666667 |

*Continuation of Table S10*

| Chromosome | Position | Par_N16 genotype | Allele frequency highlands | Allele frequency lowlands |
|---|---|---|---|---|
| 2 | 110572408 | AA | 0.1875 | 0.5588235294 |
| 1 | 209866589 | AA | 0.1315789474 | 0.8055555556 |
| 1 | 236296165 | TT | 0.6666666667 | 1 |
| 1 | 22382496 | GG | 0.9666666667 | 0.5 |
| 9 | 137157650 | CC | 0.3333333333 | 0.9333333333 |
| 1 | 174024276 | CC | 0.5714285714 | 1 |
| 1 | 44544857 | AA | 0.7142857143 | 0.1363636364 |
| 2 | 201079813 | GG | 0.7142857143 | 0.1363636364 |
| 10 | 9897331 | AA | 1 | 0.6363636364 |
| 3 | 191200376 | TT | 0.4166666667 | 0.6363636364 |
| 2 | 64929044 | GG | 0.5588235294 | 1 |
| 5 | 30486864 | AA | 0.5588235294 | 0.5 |

**Table S11.** Allele frequency of highland and lowland South American landraces at sites intersected with Par_N16 from 390 adaptive SNPs.

| Chromosome | Position | Par_N16 genotype | Allele frequency highlands | Allele frequency lowlands |
|---|---|---|---|---|
| 1 | 128373 | CC | 0.1785714286 | 0.9 |
| 10 | 1283371 | GG | 0.1 | 1 |
| 4 | 222017566 | GG | 0.275 | 0.95 |
| 3 | 135316387 | GG | 1 | 0.1666666667 |
| 10 | 10201013 | AA | 0.8333333333 | 0.1666666667 |
| 4 | 222017560 | AA | 0.275 | 0.95 |
| 7 | 158087268 | CC | 0.3 | 1 |
| 5 | 210219024 | AA | 0.2352941176 | 0.9285714286 |
| 2 | 196736161 | CC | 0.2352941176 | 0.7083333333 |
| 5 | 210219023 | AA | 0.2352941176 | 0.9285714286 |
| 4 | 181262048 | GG | 0.2352941176 | 0.9 |
| 4 | 222017569 | TT | 0.275 | 0.95 |
| 3 | 48980794 | GG | 0.7916666667 | 0.07142857143 |
| 3 | 210134894 | CC | 0.7941176471 | 0.08823529412 |
| 1 | 233262258 | TT | 0.6666666667 | 0.08823529412 |
| 7 | 158087229 | TT | 0.6 | 0.08823529412 |
| 4 | 116493754 | CC | 0.3 | 0.875 |
| 9 | 60412695 | AA | 1 | 0.5333333333 |
| 2 | 204422205 | GG | 0.2 | 0.9473684211 |
| 3 | 210134897 | TT | 0.7941176471 | 0.08823529412 |

**Dataset S1 (separate file).** Allelic similarity between Par_N16 and SNPs with significant $F_{ST}$ implicated as adaptive to Mesoamerican highland landraces.

**Dataset S2 (separate file).** Allelic similarity between Par_N16 and SNPs with significant $F_{ST}$ implicated as adaptive to Mesoamerican lowland landraces.

**Dataset S3 (separate file).** Allelic similarity between Par_N16 and SNPs with significant $F_{ST}$ implicated as adaptive to South American highland landraces.

**Dataset S4 (separate file).** Allelic similarity between Par_N16 and SNPs with significant $F_{ST}$ implicated as adaptive to South American lowland landraces..
